# Identification and Masking of Artefactual and Misleading Within-Host Variants in Deep-Sequencing SARS-CoV-2 Data

**DOI:** 10.64898/2026.03.13.711550

**Authors:** Klara M. Anker, Matthew Hall, Rosario Evans Pena, Steven A. Kemp, Joseph Clarke, Lele Zhao, David Bonsall, Nicholas Grayson, Matthew Bashton, The COVID-19 Genomics UK (COG-UK) Consortium, Ann Sarah Walker, Tanya Golubchik, Katrina Lythgoe

**Affiliations:** Pandemic Sciences Institute, University of Oxford, Oxford, UK; Big Data Institute, Nuffield Department of Medicine, University of Oxford, Oxford, UK; Department of Clinical Research, London School of Hygiene & Tropical Medicine, London, UK; Department of Biology, University of Oxford, Oxford, UK; Wellcome Centre for Human Genetics, Nuffield Department of Medicine, NIHR Biomedical Research Centre, University of Oxford, Oxford, UK; Hub for Biotechnology in the Built Environment, Northumbria University, Newcastle-Upon-Tyne, UK; Nuffield Department of Medicine, University of Oxford, Oxford, UK; The National Institute for Health Research Health Protection Research Unit in Healthcare Associated Infections and Antimicrobial Resistance at the University of Oxford, Oxford, UK; The National Institute for Health Research Oxford Biomedical Research Centre, University of Oxford, Oxford, UK; The Sydney Infectious Diseases Institute (Sydney ID), School of Medical Sciences, University of Sydney, Sydney, Australia

## Abstract

Deep sequencing data are increasingly used to study within-host viral diversity and to inform evolutionary inference. For SARS-CoV-2, analyses based on intra-host single-nucleotide variants (iSNVs) have been widely applied to quantify within-host diversity and infer transmission dynamics. However, these applications critically depend on the reliable identification of low-frequency variants, which remain vulnerable to systematic and technical artefacts. In this study, we show that recurrent artefactual iSNVs are common in large-scale SARS-CoV-2 sequencing data and can persist even under conservative minor allele frequency (MAF) thresholds. Using data from the UK’s Office for National Statistics COVID-19 Infection Survey, we demonstrate that such artefacts are predominantly sequencing centre-rather than protocol-specific. Each centre exhibits a modest, distinct set of recurrent artefactual variants showing little overlap with sites routinely masked at the consensus level. To address this, we developed a systematic, dataset-aware framework that uses recurrence within sequencing datasets to identify small, noise-adapted sets of artefactual iSNVs to mask. Applying this framework reduces spurious sharing of low-frequency variants between samples and qualitatively alters downstream inferences, including estimates of within-host diversity and transmission bottleneck sizes. Together, these findings highlight the importance of explicit, dataset-aware artefact control for robust inference from within-host variation, particularly as genomic studies increasingly seek to exploit sub-consensus diversity in rapidly evolving pathogens such as SARS-CoV-2.

## Introduction

The global emergence of SARS-CoV-2 and the evolution of multiple variants of concern (VOCs) has demonstrated the critical role of genomic surveillance in managing the pandemic through the use of whole-genome sequencing (WGS) (Zhou et al. 2020). WGS captures the genetic diversity of a viral population within a sample, with the consensus sequence essentially representing the most common allele at each position in the genome. It is the consensus sequence that is typically used for between-host phylogenetic analyses. Analysis of intra-host single-nucleotide variants (iSNVs), on the other hand, is typically used to track the evolution of the virus within-host, and to establish, for example, how many viral particles are transmitted at infection.

Despite its benefits, the application of WGS in tracking SARS-CoV-2 suffers from inconsistencies, likely due to varied sequencing methodologies, sample and analysis pipelines, and laboratory practices (Berno et al. 2022; Chiara et al. 2020; Mostefai et al. 2024; Stoler and Nekrutenko 2021). Here, we refer to observed viral variants that appear in sequencing data but are not present in sampled virus populations as “artefacts”. At the consensus level, known artefacts, which were first noticed as they appeared as common homoplasies on phylogenies (De Maio et al. 2020; Hunt et al. 2024), are often primer-(Ellegaard et al. 2025) or lab-specific (Turakhia et al. 2020), but could have various causes, including sequence alignment errors (Ellegaard et al. 2025) and changes introduced into the genome due to sample storage and preparation.

Additionally, artefacts have been observed at the within-host level. When examining within-host diversity, it is typical to exclude within-host variants with a minor allele frequency (MAF) of less than 2-5% (K. A. Lythgoe et al. 2021; McCrone et al. 2018; Poon et al. 2016). Nevertheless, variants that are generally present at low frequencies within hosts but recur across a large proportion of samples have been reported (K. A. Lythgoe et al. 2021; Martin and Koelle 2021; Mostefai et al. 2024; Tonkin-Hill et al. 2021). It is unlikely that these highly-shared iSNVs represent heritable genetic variation, given the short duration of infection and the typically tight transmission bottlenecks reported for SARS-CoV-2, and we therefore refer to them as artefactual iSNVs. (Li et al. 2022; K. A. Lythgoe et al. 2021; McCrone and Lauring 2016). Crucially, failing to mask positions in the genome where artefactual iSNVs are common can lead to erroneous conclusions about within-host genetic diversity, evolution, and transmission (K. A. Lythgoe et al. 2021; Martin and Koelle 2021).

Although the presence of artefactual iSNVs can be reduced through careful experimental design and replicate sampling (Bendall et al. 2025; Roder et al. 2023; Walter et al. 2023), the vast majority of SARS-CoV-2 sequences were generated primarily to obtain consensus genomes for phylogenetic surveillance and to track the emergence of antigenically distinct variants of concern. Examining the within-host diversity of these expensive to generate sequenced samples will allow information from this massive collection of sequences to be maximised, but to do so it is essential to develop approaches that allow quality data to be extracted while discarding artefacts.

To gain a clearer picture of the prevalence of artefactual iSNVs in SARS-CoV-2 data we analysed more than 123,000 whole-genome sequences collected as part of the United Kingdom’s Office for National Statistics Coronavirus (COVID-19) Infection Survey (ONS-CIS) (Katrina A. Lythgoe et al. 2023; Pouwels et al. 2021). We investigated different strategies for identifying and masking iSNVs that recur across a large proportion of samples. Regardless of the strategy used, we observed a large number of highly-shared artefactual iSNVs in the data, many of which were sequencing centre-specific. We demonstrate that failing to mask artefactual sites can lead to erroneous conclusions in downstream analyses, including considerably overestimating levels of within-host diversity and the size of the transmission bottleneck. We therefore advocate that all studies that utilise SARS-CoV-2 within-host diversity should make efforts to identify and mask highly-shared iSNVs and should carefully account for both noise and technical artefacts inherent to viral deep-sequencing data.

## Results

### Sequencing centre differences in iSNV abundance and sharing among samples

The ONS-CIS initially ran from the 27th April 2020 to the 13th March 2023, before being paused and then closed. We identified a total of 123,233 high-quality sequences from samples that were collected as part of the ONS-CIS, which met the criteria of a read depth ≥10x across ≥50% of positions in the genome. These sequences were generated by several laboratories throughout the course of the study using either a bait-capture approach (ve-SEQ) (Bonsall et al. 2020), or the ARTIC protocol with primer versions 3, 4, or 4.1 (Quick et al. 2016) (**Table 1, Supplementary Figure 1**).

**Table 1:** Sequencing laboratories and protocols. Samples were initially handled at different originating laboratories and in some cases sent to other laboratories for sequencing as indicated by the Sequencing lab column. The short codes refer to the originating laboratories and is the COG-UK prefixes found in the sequence names for each sample. The Sequencing centre-protocol column specifies the categorisation of samples used throughout this paper, based on the sequencing laboratory and protocol-primer version used. The number of high-quality sequences (depth of ≥10 at ≥50% of genome positions) obtained from each of the combinations are shown in the last column.

| Sequencing lab | Short code | Protocol | Library preparation kit | Sequencing platform | Sequencing centre-protocol | No. of high-quality genomes |
| --- | --- | --- | --- | --- | --- | --- |
| Wellcome Sanger Institute (SANG) | MILK | ARTIC v3 | NEB Ultra II | Illumina NovaSeq | SANG ARTIC 3 | 285 |
|  | QEUH | ARTIC v3 |  |  | SANG ARTIC 3 | 172 |
|  |  | ARTIC v4.1 |  |  | SANG ARTIC 4.1 | 64010 |
|  | BRBR | ARTIC v4.1 |  |  | SANG ARTIC 4.1 | 161 |
|  | LSPA | ARTIC v4.1 |  |  | SANG ARTIC 4.1 | 2126 |
| Northumbria University | NORT | ARTIC v3 | Illumina DNA Prep (CoronaHiT) | Illumina NextSeq 550 | NORT ARTIC 3 | 8754 |
|  |  | ARTIC v4 |  |  | NORT ARTIC 4 | 25975 |
|  |  | ARTIC v4.1 |  |  | NORT ARTIC 4.1 | 11893 |
| Quadram Institute Bioscience | NORW | ARTIC v?* | Nextera (CoronaHiT) | Illumina NextSeq 500 | NORW ARTIC ? | 900 |
| Oxford Viromics, NDM, University of Oxford | OXON | ve-SEQ | SMARTer Pico v2 | Illumina NovaSeq | OXON ve-SEQ | 6608 |
| Respiratory Virus Unit, Microbiology Services Colindale, Public Health England | PHEC | ARTIC v3 | Nextera XT | Illumina NextSeq 550 | PHEC ARTIC 3 | 2349<sup>†</sup> |

\* ARTIC primer version is unknown. Based on timings, we expect v4 or v4.1 was used.
<sup>†</sup> 2073 extra samples had no information of primer version and was excluded from this analysis.

To explore intra-host diversity within this dataset we applied a range of minimum minor allele frequency (MAF) thresholds from 2% to 20% required to call iSNVs, restricting analysis to sites with at least 1000x total read coverage. Initially, we also tested alternative read depth thresholds of 10x and 100x (**Supplementary Figure 2**), however, varying this threshold had minimal effect compared to varying the MAF threshold for calling iSNVs, and we decided to use the strict, 1000x depth threshold for subsequent analysis.

Increasing the MAF threshold generally reduced the number of observed variants per sample, but even at the same MAF thresholds the number of observed iSNVs per sample differed markedly between sequencing centres (**Figure 1** and **Supplementary Figure 2**). We emphasise that some of these sequencing centres were processing extremely high volumes of samples, with a priority on generating consensus genomes, not producing data for sub-consensus analysis. Therefore, it is unsurprising that these data exhibit greater sub-consensus noise and variation than would be expected from pipelines specifically optimised for minor variant detection. We also considered the genomic positions at which iSNVs were observed across samples. As expected, the total number of positions where iSNVs were observed decreased as MAF thresholds increased, but we also found that some positions were observed in a high proportion of samples from the same sequencing centre, even when using high MAF thresholds (**Figure 1b**). While the vast majority of iSNVs were observed in fewer than 1% of samples at any sequencing centre and MAF threshold, some centres exhibited numerous iSNV positions that were highly recurrent, appearing in as many as 20-50% of samples, particularly at the lower MAF thresholds of 3-5% (**Figure 1b**, **Supplementary Figure 3b**). Highly-shared iSNVs seemed to be sequencing centre-specific, with distinct patterns in their genomic distribution across datasets. In the NORT and NORW data, highly shared iSNVs were largely concentrated within narrow genomic regions, with several positions remaining detectable even at high MAF thresholds (**Figure 1c**). In contrast, SANG exhibited a more even, periodic distribution of highly shared iSNVs along the genome. Although many recurrent iSNVs were most apparent at lower MAF thresholds, others remained highly prevalent even when MAF thresholds were increased. Notably, clusters of highly shared variants were consistently observed around positions ∼18,600 and ∼25,200 that were shared by up to half of all samples from NORT, NORW, and SANG sequencing centres, and these signals persisted even at MAF thresholds ≥10% (**Figure 1c**).

**Figure 1:**
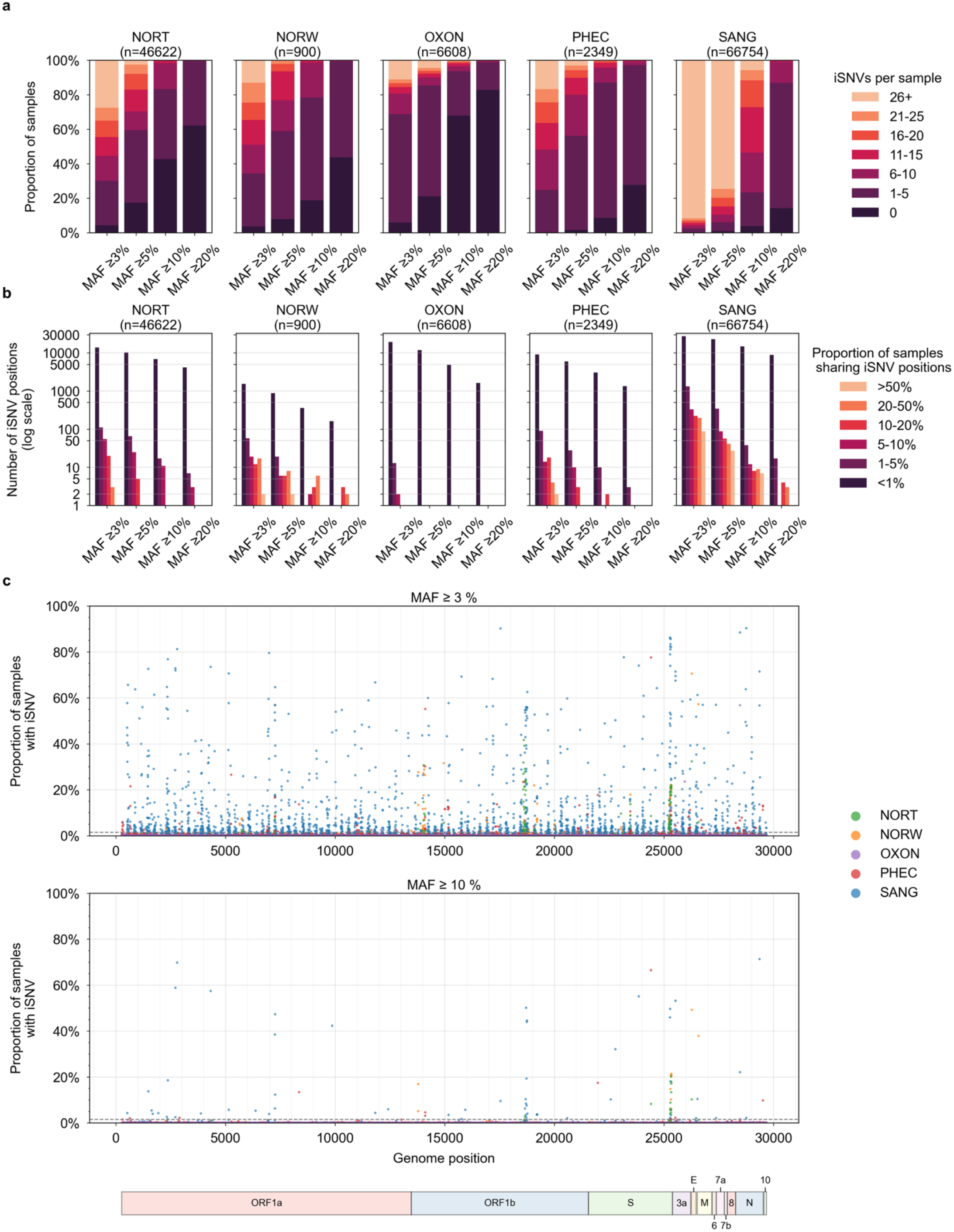
Distribution of intra-host single nucleotide variants (iSNVs) across samples processed at multiple sequencing centres, demonstrating widespread occurrence and high prevalence. **a.** Stacked bar plots showing the proportion of samples carrying different numbers of iSNVs per sample when using various MAF thresholds (3%, 5%, 10% or 20%) to call the iSNVs. Bars are colour-coded by iSNV counts. At lower MAF thresholds, samples have generally more iSNVs, whereas at higher thresholds, most samples contain few or zero iSNVs. **b.** Log10 scaled bar plots show, for different MAF thresholds, the number of distinct genomic positions that contain iSNVs in various proportions of samples. Bars are coloured by the percentage of samples sharing a given number of positions as shown on the y-axis. Most iSNV positions are shared in only a low proportion of samples, but in some sequencing centres, some positions are shared by more than half of the samples, even at higher MAF thresholds. **c.** Genome-wide map of iSNV prevalence. Each dot represents a nucleotide position (x-axis) and the percentage of samples (y-axis) that contain an iSNV at this position, coloured by sequencing centre. The top panel shows iSNVs at MAF ≥3% and the bottom panel shows iSNVs at MAF ≥10%. A grey, dashed line indicates a 1.5% proportion of samples. A schematic on the bottom shows the SARS-CoV-2 coding regions relative to the genomic positions.

### Choosing appropriate thresholds to discriminate artefactual iSNVs from true iSNVs

The observation of highly-shared, sequencing-centre specific iSNVs suggests many of them are artefactual, and should therefore be discarded for downstream analyses. Here we suggest a relatively straightforward approach for identifying artefactual iSNVs based on setting appropriate thresholds for the minimum MAF needed to call iSNVs and on the proportion of samples in which a particular iSNV is observed.

To determine appropriate ‘masking’ MAF thresholds for identifying artefactual iSNVs in the absence of duplicate sequencing, we made the pragmatic assumption that the distribution of observed iSNVs (including true and artefactual) among samples is likely to be similar across sequencing centre and protocol combinations. Here, protocol refers to whether ve-SEQ or ARTIC was used, and if ARTIC was used with primer set v3, v4 or v4.1. In a previous study, using a different SARS-CoV-2 dataset and the ve-SEQ protocol (K. A. Lythgoe et al. 2021), we used a MAF threshold of 3%, which was supported by duplicate sequencing and sequencing of synthetic genomes. Since the OXON ONS-CIS samples in this study were sequenced in the same sequencing centre and using the same protocol, we continued to use the 3% threshold for these sequences, resulting in an average of approximately 10 iSNVs per sample before masking artefactual positions (**Figure 2a**, blue dotted lines). This aligns with variant numbers previously reported by us and others (Bendall et al. 2023; K. A. Lythgoe et al. 2021; Tonkin-Hill et al. 2021). We therefore used this number as a reference to identify the MAF thresholds at which the mean number of iSNVs per sample was closest to 10 for each sequencing centre-protocol combination, with resulting MAF thresholds ranging between 3-11% (**Figure 2a**).

**Figure 2:**
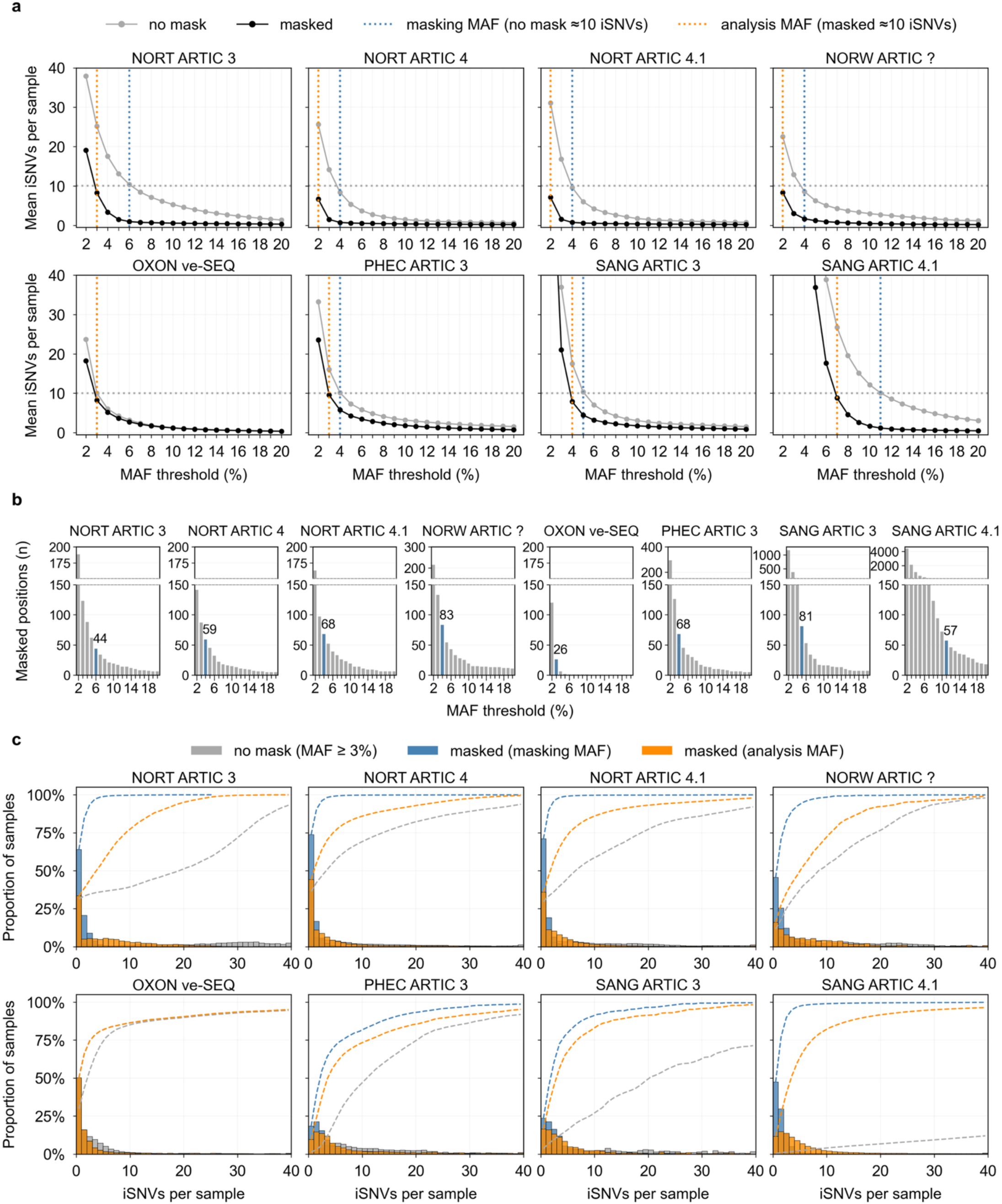
Masking of artefactual iSNVs based on varied MAF thresholds and recurrence between samples. **a**. Mean number of iSNVs per sample (y-axis) before and after masking artefactual variants for different analysis MAF thresholds (x-axis). Grey points and lines show the numbers for unmasked data, while black points and lines show the numbers after masking positions shared by more than 1.5% of samples at the ‘masking’ MAF threshold. A grey dotted line indicates the ‘reference’ mean number of iSNVs (∼10) and the ‘masking’ MAF threshold for each sequencing centre and protocol is identified as the MAF threshold where the unmasked data is closest to this reference number (blue dotted line). For example, for NORT ARTIC 4, a mean of 10 iSNVs per sample were observed before masking when using a MAF threshold of 4%, giving a masking MAF threshold of 4%. Using this MAF threshold, 59 positions had an iSNV present in more than 1.5% of samples (Figure 2b), and hence these were masked. After masking, using a 2% MAF threshold was sufficient to bring the mean number of iSNVs per sample to below 10, giving us the analysis MAF threshold (orange dotted lines). **b**. Number of genomic positions masked (y-axis) when using various masking MAF thresholds for masking iSNVs that are shared among more than 1.5% of samples. Blue bars highlight the MAF threshold and number of masked positions used in panel (**a**). Notice that all panels share the same y-axis scale until 150 masked positions, after which the y-axis is split to show more varied numbers of masked sites at low MAF thresholds for some sequencing centres. **c**. Histograms and cumulative distribution functions (CDFs) (dashed lines) showing the proportion of samples from each sequencing centre and protocol (y-axis) with various total numbers of iSNVs (x-axis). Histogram and CDF curve colours specify iSNV counts for either unmasked data analysed at a 3% MAF threshold (no mask, grey), or for masked data analysed at the masking MAF threshold (blue), or analysis MAF threshold (orange) identified for the sequencing centre and protocol in panel (**a**). X-axes have been cut at 40 iSNVs per sample and thus the full distributions are not shown in all panels.

Using these masking MAF thresholds, we then identified specific iSNVs to mask as artefactual based on how commonly they were shared among samples (**Supplementary figure 4a)**. Such a strategy will inevitably require a balance between sensitivity and specificity since true iSNVs may also appear relatively frequently in samples, especially if they are adaptive within the host. In our previous study, we used a threshold of 1.5% of samples sharing the same iSNV position to identify and mask highly-shared iSNVs, based on their MAF distributions across samples and sequencing of synthetic genomes (K. A. Lythgoe et al. 2021). Using this same 1.5% sharing threshold, and the specific MAF thresholds for each sequencing centre-protocol combination, we identified between 26 and 83 artefactual iSNV positions per combination to mask using what we call the ‘adaptive’ masking scheme (**figure 2b and Supplementary Table 1**).

Even though this resulted in masking less than 0.3% of the SARS-CoV-2 genome, with the exception of OXON the reduction in the number of observed iSNVs per sample was dramatic (**Figure 2a**, compare the grey unmasked, and black masked lines). To avoid using overly conservative MAF thresholds driven by the presence of artefacts in a high proportion of samples, which might prevent us from identifying genuine low frequency variants, we next determined ‘analysis’ MAF thresholds for each sequencing centre-protocol combination (**Supplementary figure 4b**). We still used 10 iSNVs per sample as the reference, but now after masking, this resulted in analysis MAF thresholds ranging between 2% and 7% (**Figure 2a**, orange dotted lines). This adaptive masking and analysis approach substantially reduced noise-driven variation across centres whilst also removing suspicious highly-shared variants, and made the iSNV distributions across samples converge towards similar patterns across the datasets (**Figure 2c**).

As a sense check that we were identifying a genuine signal, we tested whether the same artefactual positions were identified using the full range of MAF (2-20%) and sharing (0.5-20%) threshold combinations. Although the resultant sets of positions to mask varied greatly in size, for each sequencing centre-protocol combination we observed the expected nesting of these sets, with the smaller sets generally contained within the larger ones, and reflected by overlap coefficients close to 1 (**Supplementary Figure 4c**). This indicates that a consistent group of recurrent artefacts was detected across thresholds. The noisier datasets tended to have lower (but still high) overlap coefficients, highlighting the difficulty of distinguishing between noise, artefacts, and genuine variants. As a final sense check, we tested the masking of the most frequently flagged positions across all possible masking schemes for each sequencing centre-protocol combination, specifically positions that appeared in >20% of all threshold combinations, calling this the ‘prevalent mask’ scheme (**Supplementary Figure 5**). Apart from SANG ARTIC 4.1 this resulted in masking fewer positions, slightly more iSNVs per sample after masking, but similar analysis MAFs. The prevalent mask has the advantage of making no assumptions about the expected number of iSNVs per sample; however, by focusing only on the most frequently flagged positions, it primarily captures the strongest recurrent artefacts and may miss rarer or more context-specific artefacts.

### Highly-shared iSNVs are laboratory and protocol specific

While our suggested masking strategies resulted in comparable numbers of masked positions between sequencing centres and protocols (**Figure 2b, Supplementary Table 1**), the sets themselves were highly sequencing centre specific and did not reveal artefactual iSNVs that were consistently observed across British SARS-CoV-2 isolates (**Figure 3**). For most masking sets, approximately 50-75% of the masked positions were unique to the given sequencing centre and protocol, and only 11 out of 107 positions overlapped the previously reported list of problematic sites to mask at the consensus level (De Maio et al. 2020). At the NORT sequencing centre, many masked positions were shared across the ARTIC v3, v4 and v4.1 primers, indicating that artefacts were predominantly sequencing centre rather than primer specific. The NORT and NORW masks overlapped substantially (45 of 83 NORW masked positions were present in at least one of the NORT masks), which is likely because NORW performed over-flow sequencing for NORT when the prevalence of the virus was very high.

**Figure 3:**
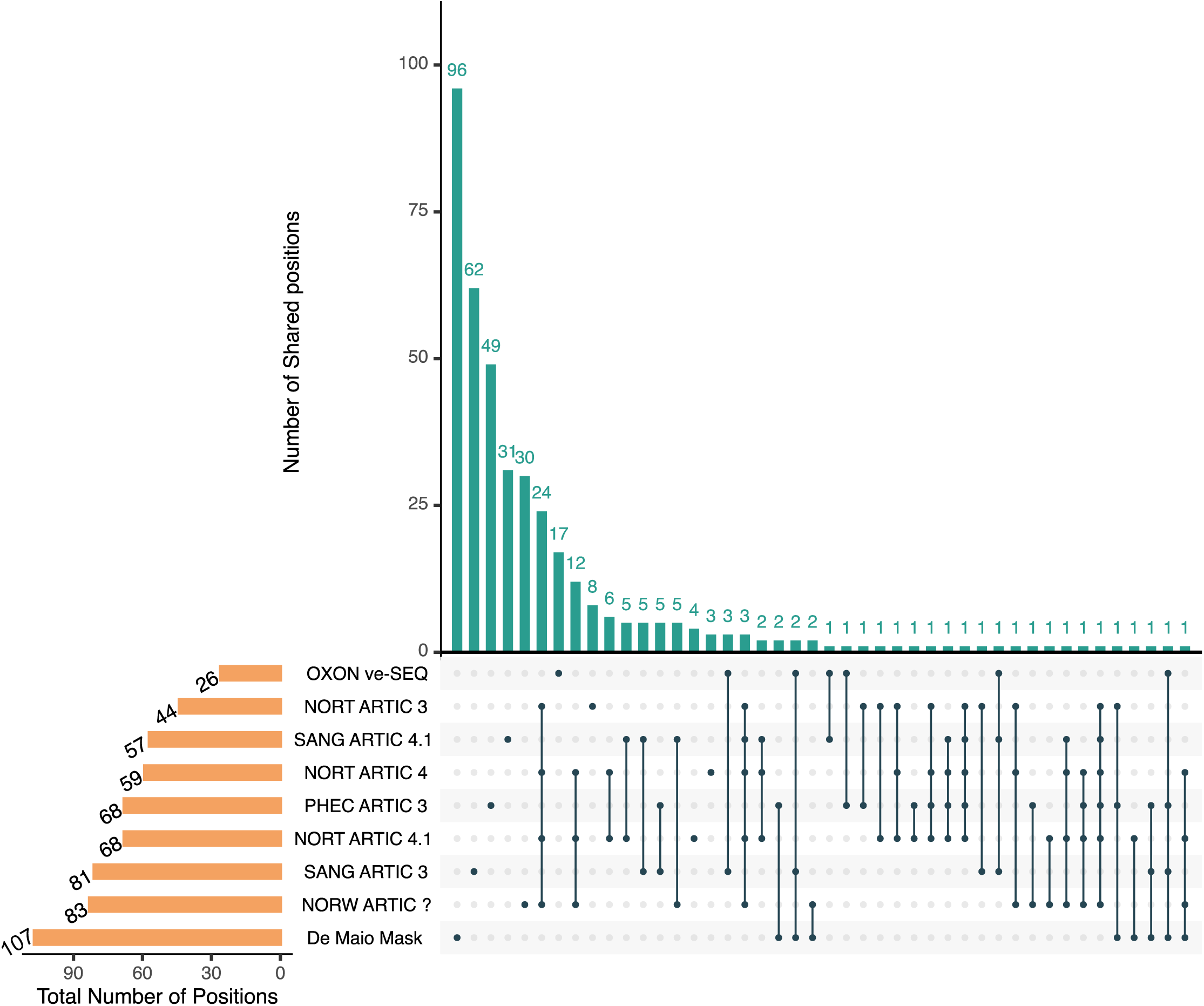
An upset plot showing intersections of masked positions using the adaptive masking strategy. Horizontal bars represent the total number of positions masked for each sequencing centre-protocol combination. The common homoplasy masking set recommended by (De Maio et al. 2020) is also shown for comparison. Points in each of the rows indicate sets of positions found within the corresponding masking set, either alone or, if connected to other dots, the number of positions shared between these mask sets given on the left. Vertical bars above the points show the number of positions in the specific intersecting sets or which are unique to one set. Most positions are sequencing centre specific.

### Considering possible sources of recurrent artefactual iSNVs

To evaluate potential sources of artefactual iSNVs, we examined strand bias, the genomic distribution of iSNVs, and their relationship to ARTIC primer-binding regions and amplicon structure. Strand bias refers to sequencing artefacts in which reads supporting a variant are unevenly distributed between forward and reverse strands and is used as an indicator of systematic errors in high-throughput sequencing data. In amplicon-based sequencing, such bias can arise from primer mismatches or uneven amplification and is therefore often used as a filter to reduce false-positive variant calls (Guo et al. 2012).

Samples from the OXON sequencing centre were excluded from this analysis, as the ve-SEQ protocol is capture-based and not subject to the primer-related strand artefacts associated with ARTIC amplicon sequencing. To test whether strand bias was associated with artefactual iSNVs, we pooled data from 100 randomly selected samples from each sequencing centre-protocol combination (see Methods). For each genomic position, we assessed whether there was a significant excess of forward or reverse reads using Fisher’s exact test and examined the overlap between significantly strand-biased positions and the corresponding mask sets. The NORT and PHEC datasets exhibited relatively few strand-biased positions and limited overlap with their respective mask sets (NORT ARTIC v3: 3/44; NORT ARTIC v4: 7/59; PHEC: 9/68), whereas the NORW dataset showed a greater number of strand-biased positions and a higher overlap with the mask set (NORW: 22/68) (**Supplementary Figure 6a-b**). In contrast, the SANG datasets contained a large number of strand-biased positions (1,325 in SANG ARTIC v3 and 930 in SANG ARTIC v4.1), with 44 of the 81 masked positions in the SANG ARTIC v3 dataset overlapping strand-biased sites (**Supplementary Figure 6c**). Strand bias therefore appears to contribute to some artefactual iSNVs in a centre- and protocol-specific manner, particularly in the SANG datasets. However, because only a subset of masked positions showed strong strand bias, it does not fully explain the artefactual iSNVs identified within each sequencing centre-protocol combination.

Given the previously observed periodic distribution of highly shared iSNVs, particularly in the SANG datasets, we next examined the distance of iSNV positions to primer-binding regions and amplicon ends. This analysis revealed peaks of highly shared iSNVs located approximately 150-170 bp from primers and amplicon ends in the SANG datasets (**Supplementary Figure 7**). Such spacing is consistent with Illumina end-of-read errors accumulating near the centres of ARTIC amplicons, as previously reported (Dong et al. 2025). SANG was the only sequencing centre to use a non-fragmentation library preparation strategy, whereas the other datasets were generated using protocols that randomly fragment amplicons prior to sequencing, which may contribute to the absence of similar patterns in those datasets.

We also assessed whether the highly shared, masked iSNVs were directly associated with primer-binding regions for each ARTIC primer version and observed only modest overlaps (**Supplementary Figure 6a-c**). To formalise this, we tested whether masked positions fell within primer binding regions more often than expected by chance using a simple hypergeometric model with the full SARS-CoV-2 genome as the background. Under this null, the expected number of overlaps is the fraction of the genome contained in primer-binding regions multiplied by the number of masked positions. Across all three primer versions, observed overlaps were close to random expectation and were not significant after correction for multiple testing (**Supplementary Table 2**), indicating that residual primer sequence or incomplete primer trimming is unlikely to be a dominant source of artefactual iSNVs.

Finally, examination of the genomic distribution of masked positions revealed that, while artefactual iSNVs were broadly dispersed across the genome for most sequencing centres (**Supplementary Figure 8a**), NORT and NORW specifically exhibited clusters of masked sites concentrated within discrete genomic intervals, some of which coincided with regions of reduced sequencing coverage (**Supplementary Figure 8b-d**). Such patterns are consistent with suboptimal amplification of specific ARTIC amplicons, which would be expected to increase stochastic error rates and promote recurrent artefactual variant calls within affected regions.

These analyses indicate that artefactual iSNVs arise through multiple, dataset-specific technical processes, none of which alone fully explains the observed patterns and thus underscores the need for adaptive, centre-specific approaches to identify and mask recurrent artefacts.

### Artefactual positions can inflate measures of within-host diversity and the transmission bottleneck size

Accurate characterisation of within-host diversity is essential for understanding within-host viral evolution and inferring transmission dynamics. However, artefactual iSNVs can inflate diversity estimates and create spurious genetic similarities between samples. To assess how such artefacts could affect downstream analyses, we compared iSNV frequencies in related and unrelated samples and estimated transmission bottleneck sizes before and after masking recurrent artefactual positions.

As a negative control, we first analysed randomly selected, non-household sample pairs from the same sequencing centres (**Figure 4a**). Prior to masking, several apparently shared variants were observed between unrelated samples, particularly at lower read-depth and MAF thresholds. In the absence of epidemiological context, such patterns could be misinterpreted as evidence of transmission. However, as read-depth thresholds were increased and MAF thresholds adapted to the overall noisiness of the data, these apparent similarities diminished. After masking highly shared variants using the adaptive masking scheme, no convincing shared within-host diversity remained, leaving only fixed consensus differences and low-frequency noise below the analysis MAF thresholds.

**Figure 4:**
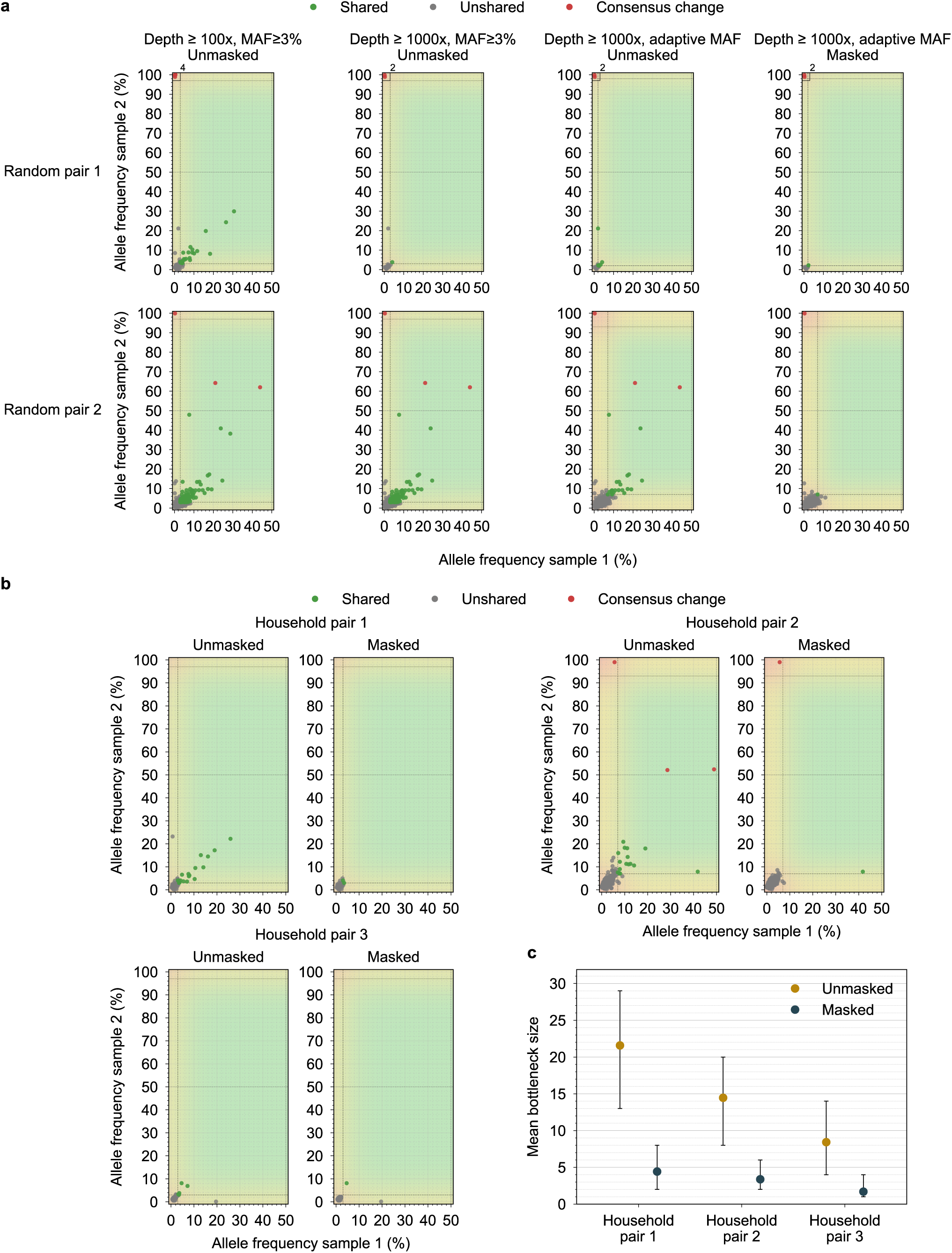
iSNV frequencies in different pairs of samples and estimated bottleneck sizes before and after masking. **a**. iSNV frequencies in two randomly paired samples processed at the same sequencing centre. The samples on the x-axis were sampled approximately one week before the samples on the y-axis, but the samples within each pair are from different households and therefore unlikely to be direct transmission pairs. Each column of plots shows variant frequencies under different criteria, using various depth and MAF thresholds to call iSNVs and with (rightmost column) or without masking highly shared, artefactual positions. Green points indicate allele frequencies of minor variants shared between the source-recipient pair, red points indicate variants where the consensus differs in the two samples, and grey points indicate minor variants that are below the analysis MAF threshold used for the given sequencing centre and protocol in either the source or recipient. The number in the top-left corner of the top row of plots gives the number of variants that are all below the MAF threshold in the first samples and close to 100% frequency in the second sample (full SNP). **b**. iSNV frequencies in three epidemiologically linked samples, where one household member (source) tested positive approximately one week before the other member (recipient). Plots on the left side show variants before masking artefacts, while plots on the right show the variants remaining after masking problematic artefact positions using the adaptive masking scheme. Here, both unmasked and masked plots show variants called under a 1000x depth and the analysis MAF threshold for the given sequencing centre and protocol. Variants are coloured as in (**a**). **c.** Estimated bottleneck size estimates for each household pair from (**b**) before and after masking artefactual sites. Gold (unmasked) and dark blue (masked) points indicate the posterior mean bottleneck estimate and error bars indicate the 95% highest posterior density intervals.

We next examined likely household transmission pairs from the ONS-CIS, focusing on households in which one individual tested positive at least one week before any other member, with subsequent household member(s) testing positive 7-10 days later, suggesting direct transmission and a clear index case. For three such pairs, we compared iSNV frequencies and fixed consensus changes between source and recipient samples before and after masking artefactual positions using the adaptive masking scheme (**Figure 4b-c**). The transmission bottleneck size was estimated using the beta-binomial method (Leonard et al. 2017) but adapted to use a Bayesian Markov Chain Monte Carlo (MCMC) approach instead of the original maximum-likelihood framework, allowing the presentation of highest posterior density intervals for the bottleneck size instead of the likelihood ratio test estimates of the original manuscript (see methods). In the first pair, unmasked data showed multiple shared iSNVs at similar frequencies in source and recipient (**Figure 4b**, Household pair 1), suggesting transmission of a genetically diverse viral population and yielding a large inferred transmission bottleneck (mean bottleneck estimate exceeding 20 virions) (**Figure 4c**). After masking, most of this apparent shared diversity disappeared, leaving only a small number of shared variants just above the analysis MAF threshold and resulting in a substantially reduced bottleneck estimate. In the second pair, the unmasked data again showed numerous shared variants across a wide frequency range, most of which were removed by masking. However, one shared iSNV and another variant that was just below the analysis MAF threshold in the source but fixed in the recipient persisted, still supporting a direct transmission link between the samples (**Figure 4b**). The third pair showed a few low-frequency shared variants, with only one remaining after masking. Across all three example pairs, masking substantially reduced the estimated bottleneck sizes, yielding smaller and more consistent estimates of approximately 2-5 virions. These narrower estimates imply that only a few virions typically initiated infection in the recipient, aligning with the expectation of tight transmission bottlenecks for SARS-CoV-2 and consistent with previous reports (Bendall et al. 2023; Hannon et al. 2022; K. A. Lythgoe et al. 2021).

## Discussion

Analyses of within-host viral variation inferred from deep sequencing data are increasingly used to study viral evolution and transmission dynamics. For SARS-CoV-2, which has been sequenced at an exceptional scale across diverse laboratories and protocols, such analyses place particularly strong demands on the reliability of low-frequency variant detection. iSNVs have been used to quantify within-host diversity, investigate putative transmission links, and estimate transmission bottleneck sizes (Bendall et al. 2023; Hannon et al. 2022; K. A. Lythgoe et al. 2021; Martin and Koelle 2021; Popa et al. 2020; Tonkin-Hill et al. 2021). These applications offer insights that are not accessible from consensus genomes alone, but they also critically rely on the assumption that the low-frequency variants predominantly reflect true biological signal rather than technical artefacts.

However, we and others have observed iSNVs that appear in the data more frequently than is biologically plausible, with some highly-shared variants persisting even when conservative MAF thresholds are applied (K. A. Lythgoe et al. 2021; Martin and Koelle 2021; Mostefai et al. 2024; Walter et al. 2023). Such recurrence raises concerns about the reliability of iSNV-based inference when systematic technical artefacts are not appropriately accounted for. Previous efforts to control likely artefactual iSNVs have largely relied on ad hoc, study-specific filtering strategies. These include restricting analyses to variants supported across technical or biological replicates (Braun et al. 2021; Grubaugh et al. 2019; McCrone and Lauring 2016; Tonkin-Hill et al. 2021), removing variants that are shared across a large proportion of samples within a dataset (K. A. Lythgoe et al. 2021), and excluding sites flagged as problematic in consensus-level analyses aimed at identifying lab-associated or recurrent error in global datasets (De Maio et al. 2020; Turakhia et al. 2020). While these approaches can remove well-characterised error modes, they are typically applied as fixed rules and do not explicitly account for variation in noise structure across datasets.

Here, we developed a systematic and dataset-aware framework for identifying and masking recurrent artefactual iSNVs. Rather than assuming a universal frequency threshold or relying on predefined problematic-site lists, our approach exploits recurrence patterns observed across samples within each sequencing dataset to identify variants that are unlikely to reflect genuine within-host variation. The framework yields small, targeted sets of positions to mask, and is designed to be scalable to diverse, large-scale sequencing datasets, while explicitly accommodating heterogeneity in noise across different sequencing centres and sample processing protocols.

Applying this framework to data from the ONS-CIS showed that recurrent artefacts are predominantly sequencing centre-specific rather than protocol-specific. Each sequencing centre-protocol combination exhibited a modest but distinct set of recurrent artefactual variants, typically numbering only a few dozen sites, with limited overlap between centres and minimal overlap with widely used consensus-level problematic-site lists (De Maio et al. 2020). This pattern indicates that systematic artefacts arise primarily from local laboratory- or batch-associated processes rather than from universal properties of sequencing protocols. Similar centre-specific structure has been observed in another large-scale analysis of public SARS-CoV-2 NGS libraries, where data-driven approaches, including dimensionality reduction, were used to distinguish systematic artefacts from putative within-host mutations (Mostefai et al. 2024). More generally, while model-based variant quality frameworks integrating multiple sources of technical bias are widely used in human genomics (e.g. GATK (Van der Auwera et al. 2013)), analogous approaches tailored to viral deep-sequencing data remain largely unexplored.

The identification and masking of artefactual iSNVs remains inherently constrained by trade-offs between sensitivity and specificity that apply to within-host analyses more generally. Even under stringent quality assurance, low-frequency artefacts can persist, and the distinction between biological signal and noise depends on how within-host variation is measured and summarised. In settings where genuine within-host evolution is expected, such as persistent infections in immunocompromised individuals, replicate sequencing can help validate minor variants (Ben Zvi et al. 2025), but such approaches are rarely feasible at scale, do not generalise to routine sequencing datasets, and may fail to identify very highly-shared artefacts as problematic since they may appear in both replicates. More broadly, measures of within-host diversity remain sensitive to sequencing depth, sampling variance, and detection thresholds, which can bias diversity estimates and complicate comparisons across samples or studies if not carefully considered (Lauring 2020; Zhao and Illingworth 2019).

Failure to identify and account for recurrent artefactual iSNVs can substantially distort biological inference based on within-host variation. Artefactual variants can inflate estimates of within-host diversity, generate spurious genetic similarity between unrelated samples, and bias transmission bottleneck estimates towards unrealistically large values. Shared low-frequency variants, when interpreted without appropriate within-dataset controls, may therefore be mistaken for evidence of direct transmission, even though such variants are often observed between epidemiologically unrelated samples (Walter et al. 2023). These methodological effects may contribute to the wide range of SARS-CoV-2 transmission bottleneck estimates reported across studies (Bendall et al. 2023; Hannon et al. 2022; K. A. Lythgoe et al. 2021; Popa et al. 2020), alongside genuine biological and epidemiological variation.

Overall, these findings show that recurrent artefactual iSNVs are common, often sequencing centre-specific, and capable of strongly biasing within-host and transmission-level inference if left unaddressed. Reliable use of iSNV data therefore requires systematic, dataset-aware artefact control rather than reliance on fixed thresholds or universal masking rules. As genomic surveillance increasingly seeks to exploit sub-consensus variation, transparent recurrence-aware strategies, together with comprehensive and standardised recording of laboratory and sequencing metadata in large sequence databases, will be essential to ensure that evolutionary and epidemiological conclusions reflect viral biology rather than technical noise, both for SARS-CoV-2 and for future pathogen surveillance efforts.

## Materials and Methods

### ONS COVID-19 Infection Survey

The ONS-CIS was a UK household-based surveillance study which ran between 26^th^ April 2020 and 31^st^ March 2023 before being paused and then closed (K. A. Lythgoe et al. 2021; Katrina A. Lythgoe et al. 2023). During this period, more than 225,000 households were recruited on a rolling basis from across the UK with the aim of being as representative as possible. Written consent was obtained from all participants and all consenting individuals aged two years and older from each household provided weekly nose + throat swab samples for the first four weeks after recruitment, and monthly swabs thereafter. All swabs were tested for SARS-CoV-2 by RT-PCR. Between April 2020 and November 2020, a selection of RT-PCR positive samples were sequenced each week, with some additional retrospective sequencing of stored samples. From December 2020 onwards, all samples with a Ct ≤30 were sent to one of several participating laboratories for sequencing.

### Sequencing

All samples analysed in this study were sequenced on Illumina platforms (MiSeq, NovaSeq, or NextSeq). Two major sequencing protocols were used: the ARTIC amplicon protocol (with primer versions 3, 4.0, and 4.1), with consensus FASTA sequence files generated using the ARTIC nextflow processing pipeline (Davis James et al. 2021; Ulhuq et al. 2023), or ve-SEQ, an RNASeq protocol based on a quantitative targeted enrichment strategy (Bonsall et al. 2015; Bonsall et al. 2020; K. A. Lythgoe et al. 2021), with consensus sequences and BAM files produced using shiver (Wymant et al. 2018). For ARTIC datasets, library preparation methods varied by sequencing centre (**Table 1**). At Northumbria University (NORT) and Quadram Institute Bioscience (NORW), libraries were prepared using the CoronaHiT method (Baker et al. 2021). Some other samples were generated using the ARTIC LoCost protocol (Quick 2020); however, the specific library preparation workflows were not consistently recorded in the available metadata.

Eight originating laboratories participated and samples were sequenced at five different sequencing laboratories. Samples are referred to these sequencing laboratories throughout this paper according to the short codes: NORT, NORW, OXON, PHEC, and SANG (**Table 1**).

### Bioinformatic processing

The processing of OXON sequence data (ve-SEQ protocol) has been previously described (K. A. Lythgoe et al. 2021). Briefly, sequences were pre-processed by downloading FASTQ files from CLIMB-COVID (Nicholls et al. 2021) and classifying them using Kraken 2 (Wood et al. 2019) with a custom database including the human genome (GRCh38) and the full bacterial/viral RefSeq set downloaded in May 2020. Human and bacterial sequences were filtered out using Castanet’s filter_keep_reads.py (Goh et al. 2019). The remaining viral and unclassified reads were trimmed to remove primers and adapters using Trimmomatic (Bolger et al. 2014) v0.36. These reads were mapped to the SARS-CoV-2 RefSeq genome (Wuhan-Hu-1 isolate MN908947.3) using shiver_map_reads.sh from the shiver (Wymant et al. 2018) v1.5.7 package, and smalt (https://github.com/rcallahan/smalt) as the read mapper. Shiver provides base frequency (BaseFreq) files, which are .csv files containing the frequencies of A, C, G, T nucleotides, gaps, and ambiguities for each position in the genome. All genomes were already aligned to the Wuhan-Hu-1 following processing with shiver, so no further aligning was required. Consensus sequences were constructed using an in-house python script which takes each BaseFreq and calculates the most common nucleotide or gap or ‘N’ at that position using a 60% majority and a read depth of 5x. If there were fewer than 5 reads at a position, ‘N’ was called. Lineages were determined using Pangolin v4.3.132 (O’Toole et al. 2021).

For all other sequencing centres, BAM files were downloaded from CLIMB as provided by the sequencing centres and processed with shiver to produce BaseFreq files.

Only samples that passed a genome coverage quality filter of minimum 10 non-gap reads across ≥50% of the genome (defined as ≥14,000 bases) were included in our analyses.

### Identifying intrahost Single Nucleotide Variants (iSNVs)

We used a custom Python script to identify iSNVs within each SARS-CoV-2 genome, excluding positions 1-265 at the 5’ end and 29675-29903 at the 3’ end, the latter encompassing the poly-A tail. An iSNV was called if the frequency of the most common minor allele at a position in the genome was above a specific threshold (ranging between 2%-20%), provided there was a total sequencing depth of at least 1000x (excluding gaps) at that position. Positions with a gap or ambiguity as the majority allele were excluded from analysis and minor allele gaps were not considered as iSNVs. Initially, we also analysed iSNVs called at positions with minimum depth thresholds of 10x or 100x total reads under the various MAF thresholds to compare with the stricter, 1000x, depth threshold.

For each sequencing centre, the proportion of samples with an iSNV at each genomic position was determined and the data was further subdivided by sequencing protocol (ARTIC primer version) if multiple protocols were used by that centre. Minor allele frequency (MAF) thresholds for calling iSNVs ranged from 2% to 20% by increments of 1%.

### Identification of highly shared iSNVs and potential masking sets

To consider both sequencing noise and artefactual, highly shared iSNVs in the data, we analysed the distribution and sharedness of iSNVs across genomic positions using various MAF thresholds and thresholds for the allowed sharedness of iSNVs among samples from the same sequencing centre and protocol. Specifically, for each MAF threshold between 2%-20% (in 1% increments) we analysed the total number of iSNVs per sample as well as the total number of samples from the same sequencing centre and protocol with an iSNV at each genomic position.

To get schemes of possible masking sets of artefactual iSNVs among samples from the same sequencing centre and protocol, we varied thresholds of allowed sharedness of iSNVs among samples from >0.5% to >20% in 0.5% increments given the varied MAF thresholds for calling iSNVs. For each combination of MAF and sharedness thresholds, we identified a set of positions where iSNVs were called “too frequently” and thus considered these artefact positions to mask. E.g. if the MAF threshold was set to 2% and the sharedness threshold at 1.0%, every position where more than 1% of samples from a given sequencing centre/protocol category had a minor variant with a frequency of 2% or more was added to the masking set. This resulted in up to 760 different possible masking schemes for each sequencing centre and protocol.

### Choosing masking sets and MAF thresholds for true versus artefactual iSNVs

To determine a set of masking positions for each sequencing centre and protocol that considered both noise and sequencing artefacts while preserving the majority of true iSNVs, we constructed adaptive masks using the OXON dataset as a reference and a 1.5% threshold for the proportion of samples sharing the same iSNV positions as previously used in (K. A. Lythgoe et al. 2021). Using the mean number of iSNVs per sample as a metric for noise in the data, we identified the MAF thresholds at which data from each sequencing centre and protocol had a mean number of iSNVs per sample closest to the number observed for samples processed with the OXON protocol (∼10 iSNVs). This MAF threshold was then used for masking positions where >1.5% of samples from the given sequencing centre and protocol had an iSNV at or above the threshold. Post masking, samples were again analysed for their average number of iSNVs and an “analysis” MAF threshold for each dataset was proposed where the masked data were now closest to having the reference average number of iSNVs per sample.

As an alternative masking strategy, sets were also compiled from the most commonly masked positions across all tested MAF and sharedness threshold combinations for each sequencing centre and protocol. For these sets, every position that was present in >20% of possible masking schemes for a sequencing centre-protocol combination was considered artefactual and masked. Using this scheme (the prevalent mask), MAF thresholds for subsequent analysis could again be determined with the OXON data as a reference, aiming for an average number of iSNVs per sample close to 10.

### Analysing strand bias across a subset of samples from each sequencing centre

We analysed strand bias across a random subset of 100 samples per sequencing centre-protocol combination to identify positions where the distribution of minor variant reads between forward and reverse strands was systematically biased. Samples from the OXON sequencing centre were excluded because the ve-SEQ protocol (capture-based) is not prone to the same primer-related strand artefacts as ARTIC amplicon sequencing. Variant calling was performed with LoFreq (Wilm et al. 2012) (v2.1.5) against the SARS-CoV-2 reference genome (MN908947.3) with default post-filters disabled (--no-default-filter). We parsed the VCF output per sample, extracting strand-specific counts from the INFO column DP4 (ref-forward, ref-reverse, alt-forward, alt-reverse) and allele frequencies from AF.

For each sequencing centre-protocol combination, at each genomic position between 265–29,674, we pooled strand-specific counts across the subsampled set, provided at least 10 samples contributed data and the pooled depth was ≥1000 reads. The major versus minor allele was defined per sample using AF (if AF>0.5 the alternative allele, ALT, is major, otherwise the reference, REF, is major). When multiple ALT<0.5 were present, the one with the most reads was used as the minor allele. We then tested a 2x2 table (major/minor x forward/reverse) using a two-sided Fisher’s exact test. Positions were excluded if underpowered: <10 contributing samples, pooled depth <1000 reads, any zero cell or expected cell count <5 in the 2x2 table, or sensitive to a small pseudo-count (adding 0.5 to all cells changed the p-value by >0.1). Raw p-values were adjusted across positions using the Benjamini-Hochberg procedure and finally positions with q<0.01 were designated as strand-biased for that sequencing centre-protocol combination and used in downstream comparisons with other mask sets.

### Comparison between masking sets and known problematic positions

To characterise and analyse possible explanations for the sets of masked iSNVs in each sequencing centre, we compared each masking set to each other and to sets of potentially problematic sites. These included (1) known problematic positions (De Maio Mask and De Maio Caution) (De Maio et al. 2020), (2) primer binding positions from ARTIC primer schemes v3, v4, and v4.1, and (3) positions with significant strand bias identified across subsets of samples from each sequencing centre. Overlaps and unique positions among these sets were visualised using UpSet plots generated with the UpSetR R package (Conway et al. 2017).

### Test of enrichment of masked sites in primer-binding regions

For each ARTIC primer version (v3, v4, v4.1) we tested whether masked sites occurred within primer-binding regions more often than expected by chance. For each version, we formed the union of unique masked genomic positions across masking sets using that primer scheme (e.g. NORT ARTIC 3, PHEC ARTIC 3, SANG ARTIC 3). Primer-binding coordinates were taken from the corresponding ARTIC scheme (downloaded from https://github.com/artic-network/primer-schemes/tree/master/nCoV-2019). Using the full SARS-CoV-2 genome as background (N = 29,903), we let K be the number of primer positions, M the number of masked positions in the union, and x the observed overlap. Under a null of uniform placement without replacement, *X* ∼ *Hypergeom* (*N*, *K*, *M*). One-sided enrichment p-values were computed as *p* = Pr[*X* ≥ *x*], using phyper in R. We also reported primer coverage *K*/*N*, the expected overlaps *M* × *K*/*N*, and fold enrichment (observed/expected). Because three tests were prespecified (one per primer version), p-values were Benjamini–Hochberg adjusted and reported as q-values.

### Bottleneck size calculation

The minor allele frequencies from three likely transmission pairs were analysed using a modified version of the exact beta-binomial method of (Leonard et al. 2017). In this study, sites were considered as potential iSNVs only if the depth at that position was ≧1000 reads in the source individual, and the iSNV frequency at that position was above the analysis MAF threshold for the source’s sequencing centre.

A Bayesian Markov Chain Monte Carlo (MCMC) approach was used instead of the original maximum-likelihood framework, allowing the presentation of highest posterior density intervals for the bottleneck size instead of the likelihood ratio test estimates of the original manuscript. The MCMC approach requires a prior distribution for the size of the bottleneck, which in this case was a (shifted) geometric distribution with a success probability of 0.5, resulting in a mean of 2 virions. Each MCMC chain was run for 50,000 iterations, with the first 5,000 discarded as burn-in. This process was repeated twice for each pair, first without masking any genome positions and then with masking using the adaptive mask scheme. If the source and recipients of the pair came from different sequencing centres, all positions from both mask lists were masked. The variant calling threshold in the recipient was set to the analysis threshold for the recipient’s sequencing centre.

## Data and Code Availability

All custom code written and described in this manuscript are available at https://github.com/lythgoe-lab/SC2_artefacts. All sequences are publicly available via the COG-UK project on the ENA, (https://www.ebi.ac.uk/ena/browser/view/PRJEB37886).

## Notes

This work contains statistical data from ONS which is Crown Copyright. The use of the ONS statistical data in this work does not imply the endorsement of the ONS in relation to the interpretation or analysis of the statistical data. This work uses research datasets which may not exactly reproduce National Statistics aggregates.

## Author Contribution Statement

T.G., M.H. and K.L. conceptualised the study. K.M.A., T.G., D.B., N.G., MB., M.H., R.E.P., K.L., and LZ developed the methodology. K.M.A., S.A.K., J.C., M.H. and T.G. performed formal analysis. M.H., A.S.W. and K.L. provided supervision and oversight. K.M.A, S.A.K. and K.L. wrote the original draft of the manuscript, and all authors reviewed and edited the manuscript.

