## Supplemental Material for "Identification and Masking of Artefactual and Misleading Within-Host Variants in Deep-Sequencing SARS-CoV-2 Data"

Supplementary Tables and figures

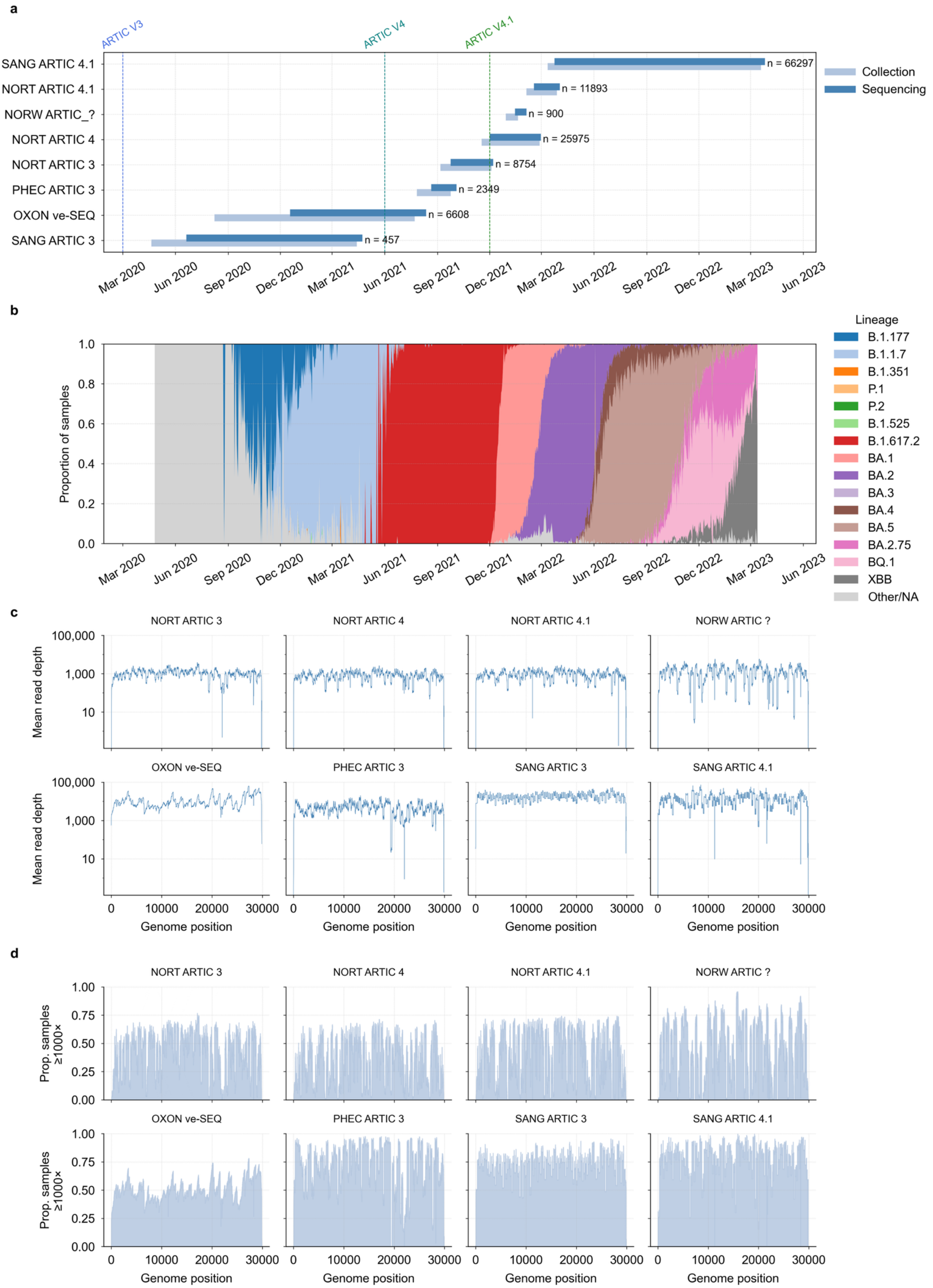
a
ARTIC V3
ARTIC V4
ARTIC V4.1
SANG ARTIC 4.1
NORT ARTIC 4.1
NORW ARTIC_?
NORT ARTIC 4
NORT ARTIC 3
PHEC ARTIC 3
OXON ve-SEQ
SANG ARTIC 3
n = 66297
n = 11893
n = 900
n = 25975
n = 8754
n = 2349
n = 6608
n = 457
Collection
Sequencing
Mar 2020
Jun 2020
Sep 2020
Dec 2020
Mar 2021
Jun 2021
Sep 2021
Dec 2021
Mar 2022
Jun 2022
Sep 2022
Dec 2022
Mar 2023
Jun 2023
b
Proportion of samples
Lineage
B.1.177
B.1.1.7
B.1.351
P.1
P.2
B.1.525
B.1.617.2
BA.1
BA.2
BA.3
BA.4
BA.5
BA.2.75
BQ.1
XBB
Other/NA
c
NORT ARTIC 3
NORT ARTIC 4
NORT ARTIC 4.1
NORW ARTIC ?
OXON ve-SEQ
PHEC ARTIC 3
SANG ARTIC 3
SANG ARTIC 4.1
Mean read depth
Genome position
d
Prop. samples ≥1000×
Genome position

**Supplementary Figure 1.** Overview of sample collection and sequencing timelines, observed SARS-CoV-2 lineages and sequencing depth across all ONS-CIS genomes. **a**. Timeline of sample processing by sequencing centre and protocol combination (y-axis) across the study period (x-axis). For each track, the dark blue bar shows the range of sequencing dates and the light blue bar indicates the range of sample collection dates for those samples. The total number of samples from each sequencing centre/protocol combination used in this study (meeting quality requirements of a read depth ≥10x across ≥50% of the genome) are shown to the right of each set of bars. Vertical lines mark the release dates of relevant versions of the ARTIC primer sets. **b**. Timeline of the proportions of samples in our dataset assigned to each of the SARS-CoV-2 PANGO lineages per sample collection date (x-axis). **c.** Average read depth across the genome for all samples from a given sequencing centre using various protocols and primer sets as indicated by the labels above each plot. **d.** Proportion of samples from each sequencing centre/protocol-primer sets with ≥1000x reads across each genome position. The plots only show "high quality" samples, classified as samples with a read depth ≥10x across ≥50% of the genome. In our analysis, only high-quality samples were included and iSNVs were only called at positions reaching a depth of 1000x.

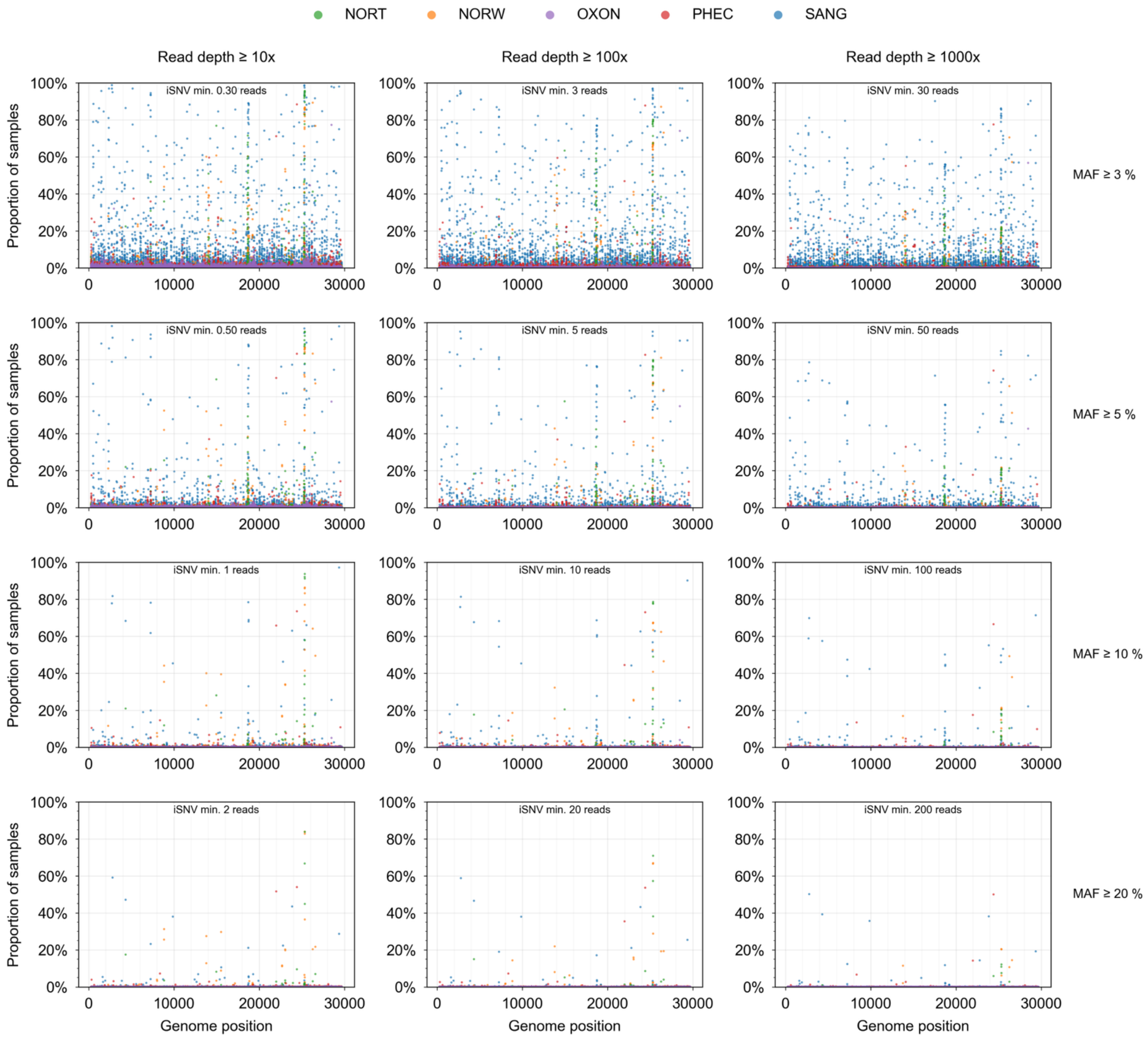

**Supplementary Figure 2**

Distribution and proportion of iSNVs given various MAF thresholds (rows) and read depth thresholds (columns) to consider a position intra-host variant. Each dot represents a nucleotide position (x-axis) and the percentage of samples (y-axis) that contain an iSNV at this position, coloured by sequencing centre (not distinguishing between different ARTIC protocols used). Labels in the top of each panel show the minimum number of reads required to call an iSNV given the read depth and MAF thresholds (thresholds <1 would in practise mean minimum 1 read to call the iSNV).

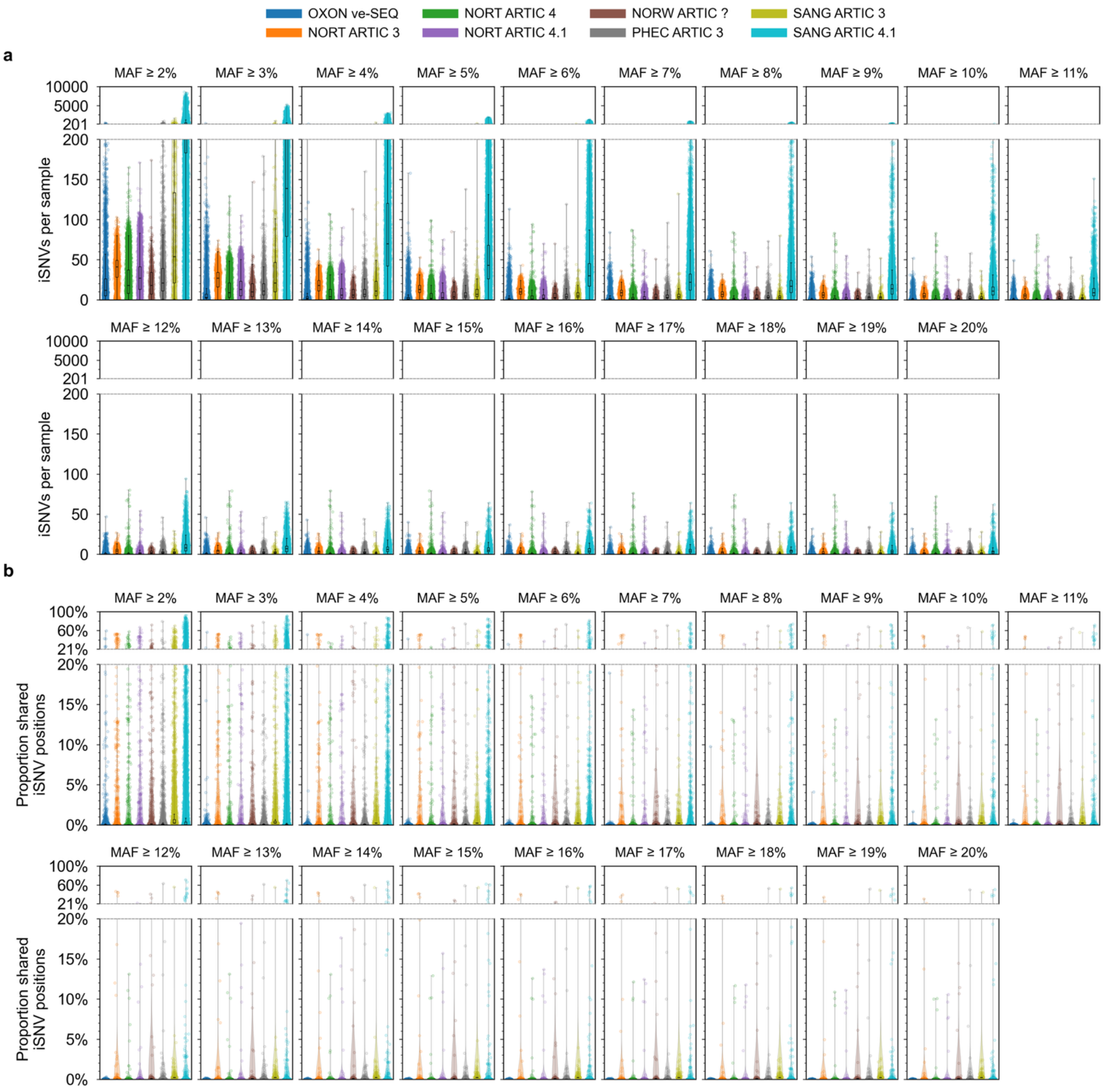


**Supplementary Figure 3.** Distribution of iSNVs per sample and shared between samples for each sequencing centre-protocol combination. **a**. Violin- and box plots of number of iSNVs per sample with each of the sequencing centre-protocol combinations shown by different colours. Each panel shows the number of iSNVs when using different MAF thresholds from 2%-20%. Y-axes are split, showing higher resolution in lower panels from 0-200 iSNVs per sample and a larger span from 201-10000 iSNVs per sample in top panels. **b**. Violin- and box plots of the proportion iSNVs that are shared between samples, with each sequencing centre-protocol combination shown by different colours. Each panel shows the proportions of shared iSNVs when using different MAF thresholds from 2%-20%. Y-axes are split, showing a higher resolution in lower panels from 0-20% shared iSNV positions and a larger span from 21-100% shared iSNVs in the top panels.

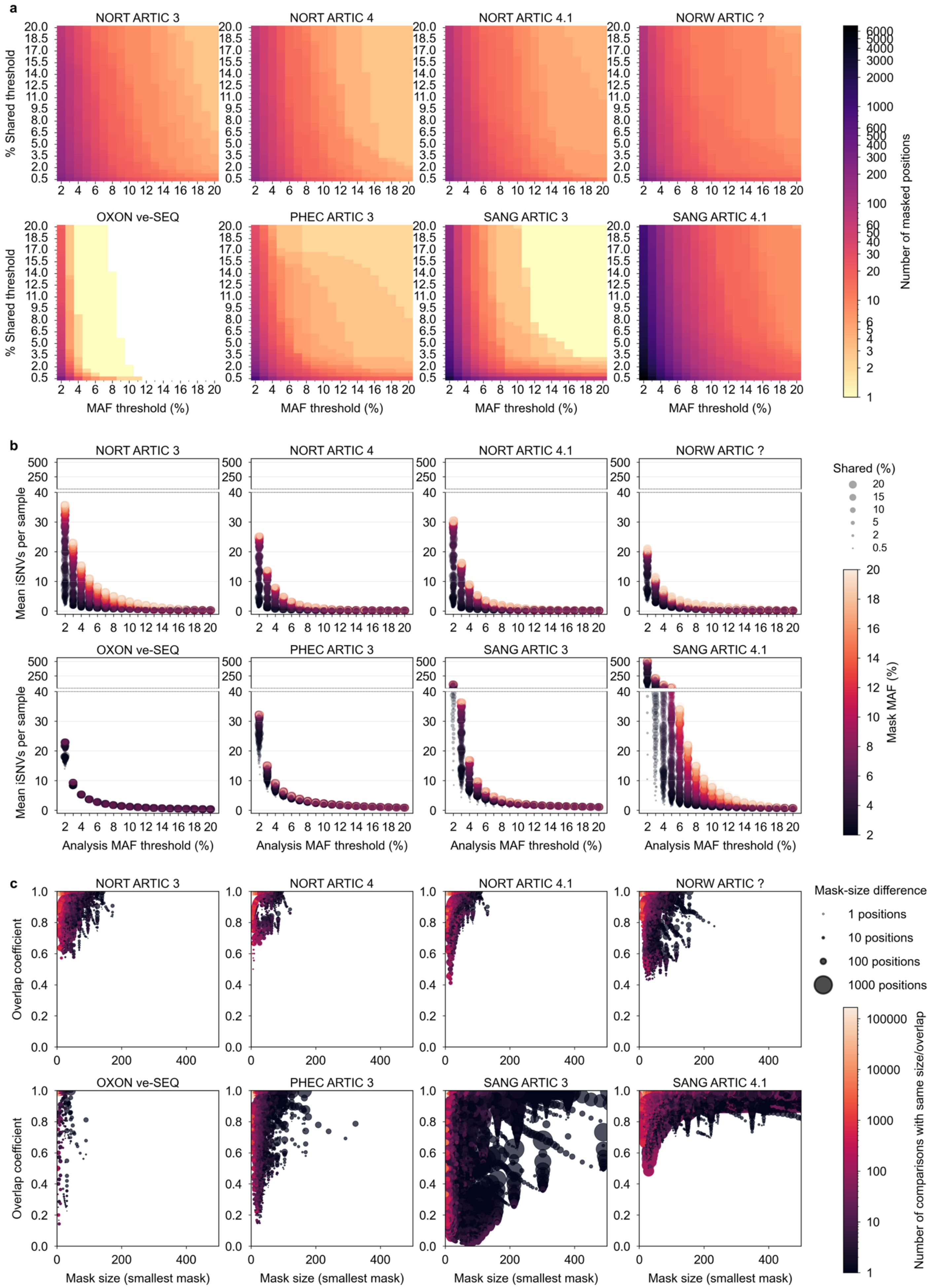

a
NORT ARTIC 3
NORT ARTIC 4
NORT ARTIC 4.1
NORW ARTIC ?
OXON ve-SEQ
PHEC ARTIC 3
SANG ARTIC 3
SANG ARTIC 4.1
% Shared threshold
20.0 18.5 17.0 15.5 14.0 12.5 11.0 9.5 8.0 6.5 5.0 3.5 2.0 0.5
2 4 6 8 10 12 14 16 18 20
MAF threshold (%)
Number of masked positions
6000 5000 4000 3000 2000 1000 600 500 400 300 200 100 60 50 40 30 20 10 6 5 4 3 2 1
b
NORT ARTIC 3
NORT ARTIC 4
NORT ARTIC 4.1
NORW ARTIC ?
OXON ve-SEQ
PHEC ARTIC 3
SANG ARTIC 3
SANG ARTIC 4.1
Mean iSNVs per sample
500 250 40 30 20 10 0
Analysis MAF threshold (%)
Shared (%)
20 15 10 5 2 0.5
Mask MAF (%)
20 18 16 14 12 10 8 6 4 2
c
NORT ARTIC 3
NORT ARTIC 4
NORT ARTIC 4.1
NORW ARTIC ?
OXON ve-SEQ
PHEC ARTIC 3
SANG ARTIC 3
SANG ARTIC 4.1
Overlap coefficient
1.0 0.8 0.6 0.4 0.2 0.0
0 200 400
Mask size (smallest mask)
Mask-size difference
1 positions
10 positions
100 positions
1000 positions
Number of comparisons with same size/overlap
100000 10000 1000 100 10 1

**Supplementary Figure 4:** Mask sizes and overlap between different masking sets of positions for each sequencing centre-protocol combination. **a.** Heatmaps showing the number of positions in the masking set for each of the combinations of MAF thresholds (x-axis) and percentages of sharedness between samples (y-axis) for each sequencing centre-protocol combination. Colours on a log scale indicates the number of positions in the masking set for each of the mask scenarios with lighter colours representing fewer positions and darker colours representing more positions. White tiles mean that no positions are in the masking set for the given MAF and sharedness threshold combination. **b.** Effect of masking with each potential mask set on the mean number of iSNVs per sample (y-axis) when analysing under various analysis MAF thresholds (x-axis) for each sequencing centre and protocol. Each point corresponds to the mean number of iSNVs per sample. The colour and size of points indicate the mask scheme with colours corresponding to the MAF threshold used for masking and sizes corresponding to the allowed percentage of sharedness under that MAF threshold. The y-axis is split in two enabling better visualisation of more extreme y-values on another scale at the lowest analysis MAF thresholds for the SANG data. **c**. Overview of the similarity between all the possible masking sets given each of the MAF and sharedness threshold combinations shown in (**a**). Bubbles indicate, for each pairwise comparison of two mask sets, the overlap coefficient given as the size of the intersection between the two sets divided by the size of the smaller set (y-axis). An overlap coefficient of 1 means the smaller of the mask sets is completely nested within the larger. The size of the smaller mask set in the comparison is indicated on the x-axis, and the size of the bubbles indicate the difference in number of positions in each of the two mask sets. A log-scale colour gradient helps highlight where many pairwise comparisons ended up at the same coordinates in the grid.

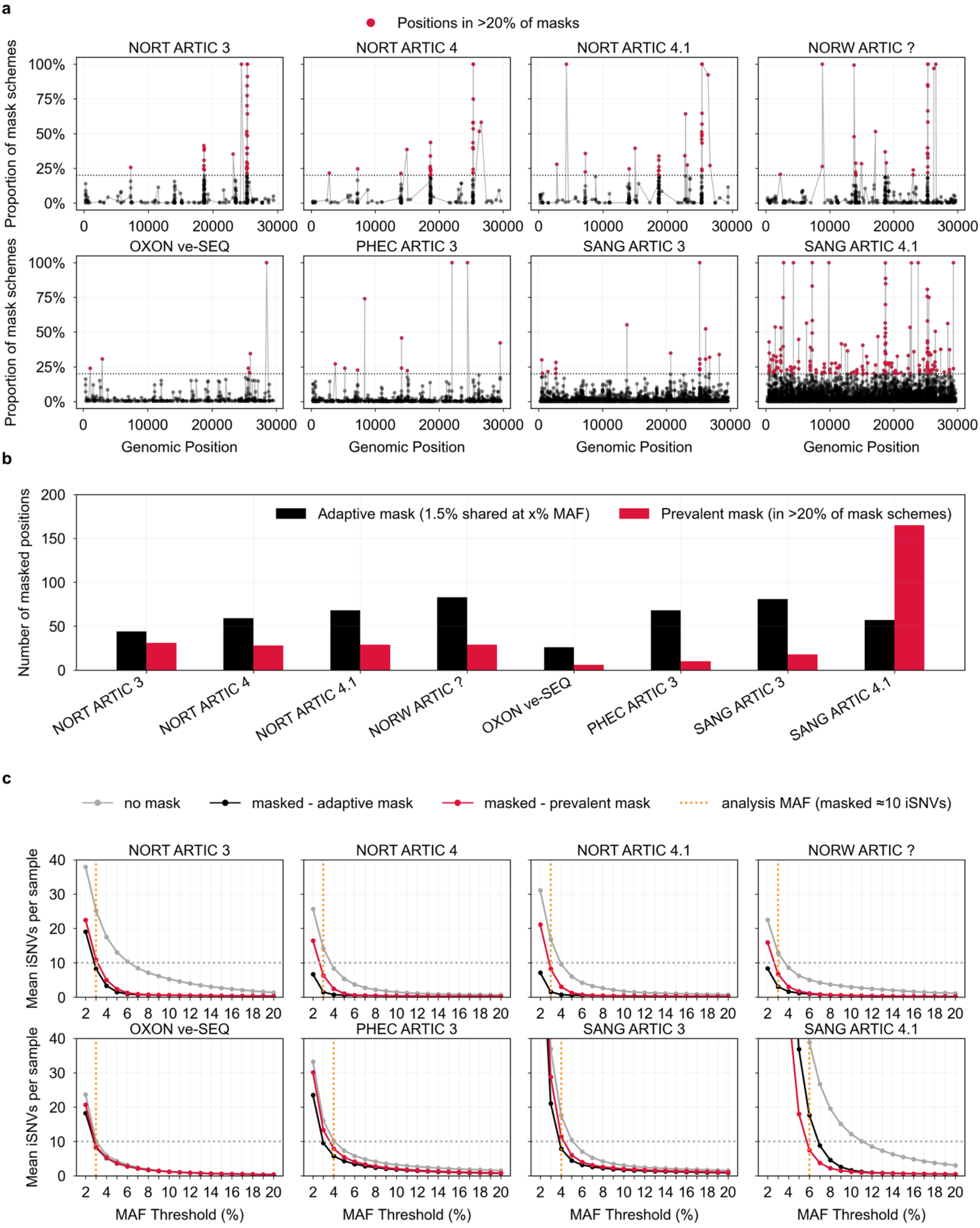


**Supplementary Figure 5:** Alternative masking strategy based on the most prevalently masked positions across all thresholds (prevalent mask). **a.** The frequency of masking different genomic positions across all the mask threshold combinations (MAF 2-20% and 0.5%-20% sharedness) for each sequencing centre and protocol. Positions are shown on the x-axis and their frequencies given as the proportion of all possible mask schemes for that sequencing centre and protocol containing the specific positions are shown on the y-axis. Dots for each masked position are connected to better show how they spread along the genome and a dotted line as well as red highlighting indicate positions prevalent in >20% of possible masks. **b.** Comparison of number of masked positions in the two suggested masking strategies. For each sequencing centre and protocol (x-axis) black bars show the number of positions masked by the adaptive masks while red bars show the number of masked positions using the prevalent strategy for the given sequencing centre and protocol. **c.** Plots showing the mean number of iSNVs per sample (y-axis) for each sequencing centre and protocol when varying the MAF threshold from 2-20% (x-axis). Grey points and lines show the numbers for unmasked data, black points and lines show the numbers after masking positions with the adaptive strategy (as in **Figure 2**), and red points and lines show the numbers after masking positions with the prevalent strategy. The horizontal blue, dotted line shows the expected average number of iSNVs per sample based on the "reference" OXON dataset and the vertical orange, dotted lines indicate the MAF thresholds (analysis MAF threshold) where the data have the expected mean number of iSNVs per sample after masking the positions seen in >20% of masks. Y-axes have been cut at 40 iSNVs per sample, and thus a few datapoints are missing for the lowest MAF thresholds for some datasets.

| Sequencing centre-protocol | Mask | Masking MAF (%) | Analysis MAF (%) | Number of masked positions |
|---|---|---|---|---|
| NORT ARTIC 3 | Adaptive (shared >1.5% of samples) | 6 | 3 | 44 |
| | Prevalent (Positions in >20% of masks) | - | 3 | 31 |
| NORT ARTIC 4 | Adaptive (shared >1.5% of samples) | 4 | 2 | 59 |
| | Prevalent (Positions in >20% of masks) | - | 3 | 28 |
| NORT ARTIC 4.1 | Adaptive (shared >1.5% of samples) | 4 | 2 | 68 |
| | Prevalent (Positions in >20% of masks) | - | 3 | 29 |
| NORW ARTIC ? | Adaptive (shared >1.5% of samples) | 4 | 2 | 83 |
| | Prevalent (Positions in >20% of masks) | - | 3 | 29 |
| OXON ve-SEQ | Adaptive (shared >1.5% of samples) | 3 | 3 | 26 |
| | Prevalent (Positions in >20% of masks) | - | 3 | 6 |
| PHEC ARTIC 3 | Adaptive (shared >1.5% of samples) | 4 | 3 | 68 |
| | Prevalent (Positions in >20% of masks) | - | 4 | 10 |
| SANG ARTIC 3 | Adaptive (shared >1.5% of samples) | 5 | 4 | 81 |
| | Prevalent (Positions in >20% of masks) | - | 4 | 18 |
| SANG ARTIC 4.1 | Adaptive (shared >1.5% of samples) | 11 | 7 | 57 |
| | Prevalent (Positions in >20% of masks) | - | 6 | 165 |

**Supplementary Table 1:** Final masks and thresholds for each dataset. Two different types of masking strategies (adaptive and prevalent) are shown along with the MAF thresholds used for masking positions and analysing iSNVs after masking. For the consensus masks no masking MAF threshold is shown, as this strategy identified positions across all tested masking MAF thresholds.

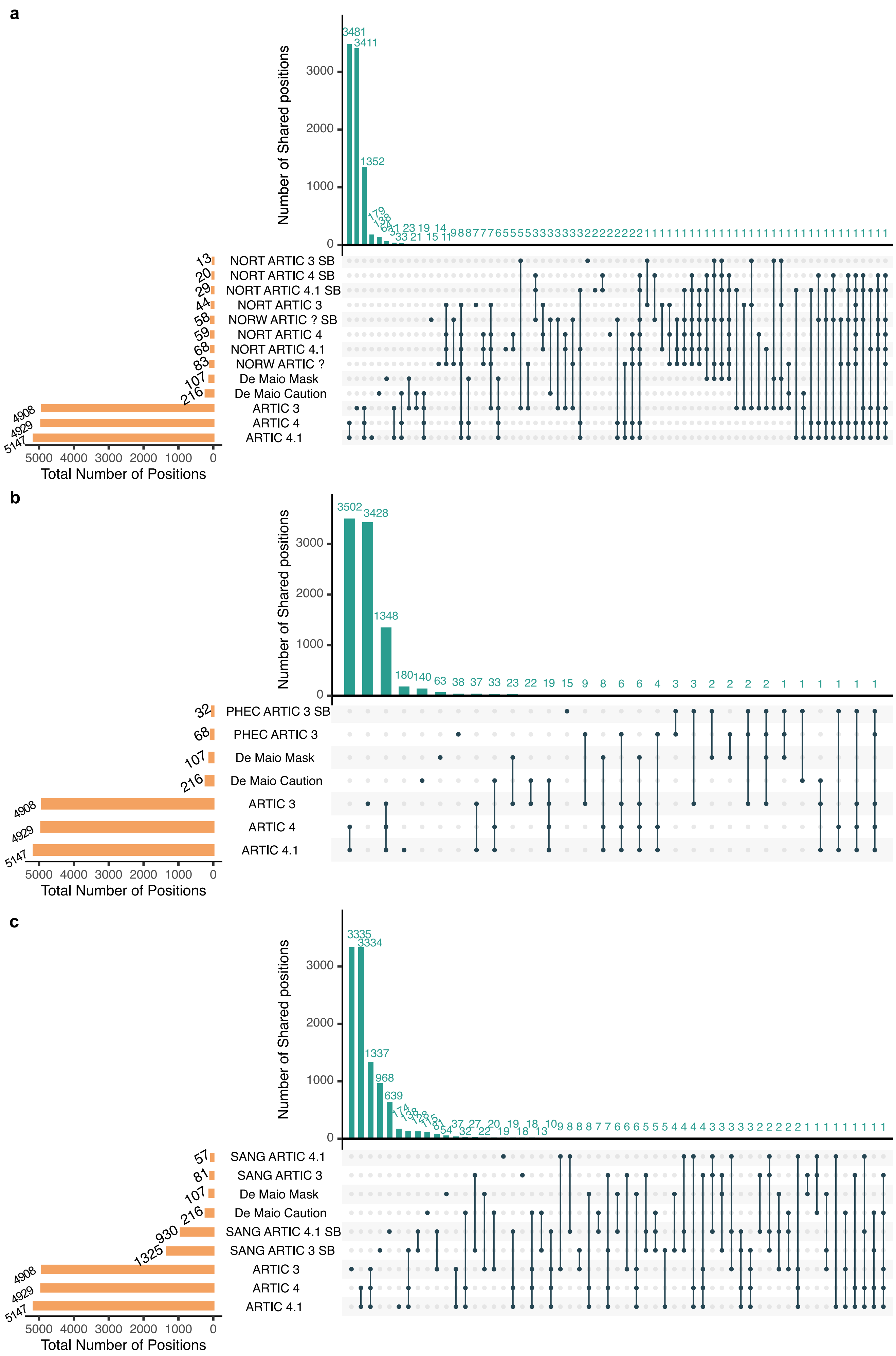

a
Number of Shared positions
3481
3411
1352
NORT ARTIC 3 SB
NORT ARTIC 4 SB
NORT ARTIC 4.1 SB
NORT ARTIC 3
NORW ARTIC ? SB
NORT ARTIC 4
NORT ARTIC 4.1
NORW ARTIC ?
De Maio Mask
De Maio Caution
ARTIC 3
ARTIC 4
ARTIC 4.1
Total Number of Positions
b
Number of Shared positions
3502
3428
1348
PHEC ARTIC 3 SB
PHEC ARTIC 3
De Maio Mask
De Maio Caution
ARTIC 3
ARTIC 4
ARTIC 4.1
Total Number of Positions
c
Number of Shared positions
3335
3334
1337
968
639
SANG ARTIC 4.1
SANG ARTIC 3
De Maio Mask
De Maio Caution
SANG ARTIC 4.1 SB
SANG ARTIC 3 SB
ARTIC 3
ARTIC 4
ARTIC 4.1
Total Number of Positions

**Supplementary Figure 6:** Upset plots showing intersections of adaptive mask sets and positions with significant strand bias (SB) across a subset of samples from **a**) NORT and NORW, **b**) PHEC, and **c**) SANG protocols. In each panel, the masked and SB positions are also compared to previously reported positions to mask or be cautious about by (De Maio et al. 2020), and positions within primer binding sites for each of the three ARTIC primer schemes used in the protocols (labelled ARTIC 3, ARTIC 4, and ARTIC 4.1. Horizontal bars represent the total number of positions in the given set. Points in each of the rows indicate sets of positions found within the corresponding set, either alone or, if connected to other dots, the number of positions shared between the sets given on the left. Vertical bars above the points show the number of positions in the specific intersecting sets or which are unique to one set.

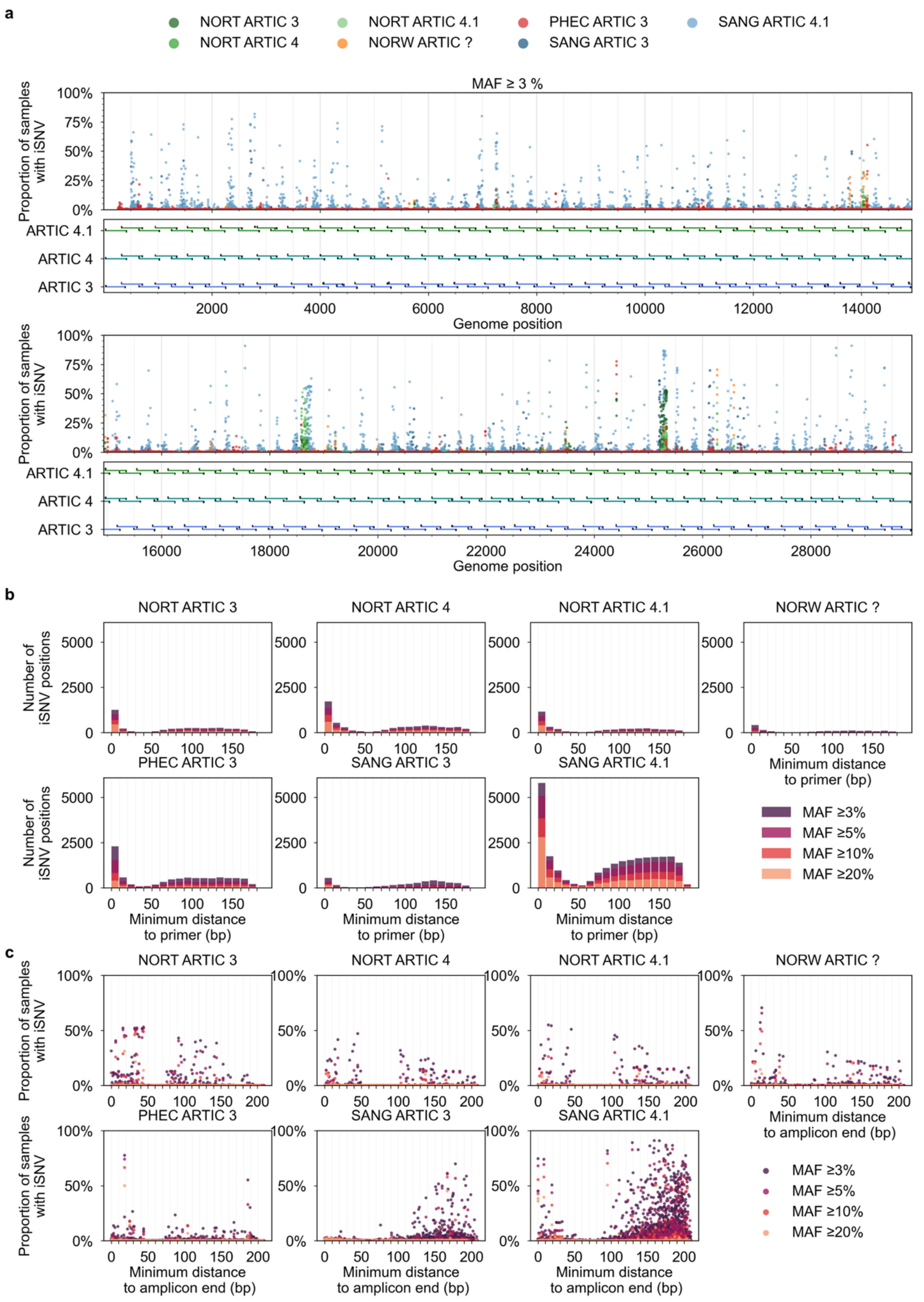


**Supplementary Figure 7**: iSNV distributions across amplicons and their distances to primers from the used ARTIC primer schemes. **a**. Genome-wide map of iSNV prevalence as in **Figure 1c**, but with lower panels showing regions covered by different amplicons for each of the three used ARTIC schemes in different colours. Small black lines, usually at the ends of each amplicon, indicate where primers (including alternative primers in each of the schemes) bind. **b**. Histograms of the number of iSNVs per sequencing-centre and protocol at various minimum distances from primer binding sites shown in bins of 10 bp distances. Due to the design of the ARTIC primer schemes, some genome positions will be covered by several amplicons; the distances shown here is the minimum distance to any primer of an amplicon covering the position. Colours indicate various MAF thresholds for identifying iSNVs. **c**. Proportion of samples with iSNVs at specific positions and the minimal distance of those positions to an amplicon end. Colours of each point indicate the MAF threshold used for calling the iSNVs.

| Primers | Background (N) | Primer positions (K) | Masked positions (M) | Observed overlap | Primer coverage fraction | Expected overlap | Fold enrichmen t | p-value | q-value |
|---|---|---|---|---|---|---|---|---|---|
| **ARTIC v3** | 29903 | 4908 | 180 | 38 | 0.164 | 29.5 | 1.29 | 0.058 | 0.086 |
| **ARTIC v4** | 29903 | 4929 | 59 | 15 | 0.165 | 9.7 | 1.54 | 0.052 | 0.086 |
| **ARTIC 4.1** | 29903 | 5147 | 145 | 29 | 0.172 | 25.0 | 1.16 | 0.215 | 0.215 |

**Supplementary Table 2:** One-sided hypergeometric test of enrichment of masked sites in ARTIC primer-binding regions. For each primer set (v3, v4, v4.1), the union of unique masked positions from datasets using that primer version was tested against the corresponding primer-binding positions using the full SARS-CoV-2 genome as background (N). K = primer binding positions; M = masked positions; Expected overlap = M×K/N; Fold enrichment = observed/expected; p-value = raw one-sided p-value; q-value = Benjamini-Hochberg (BH) adjusted across the three tests.

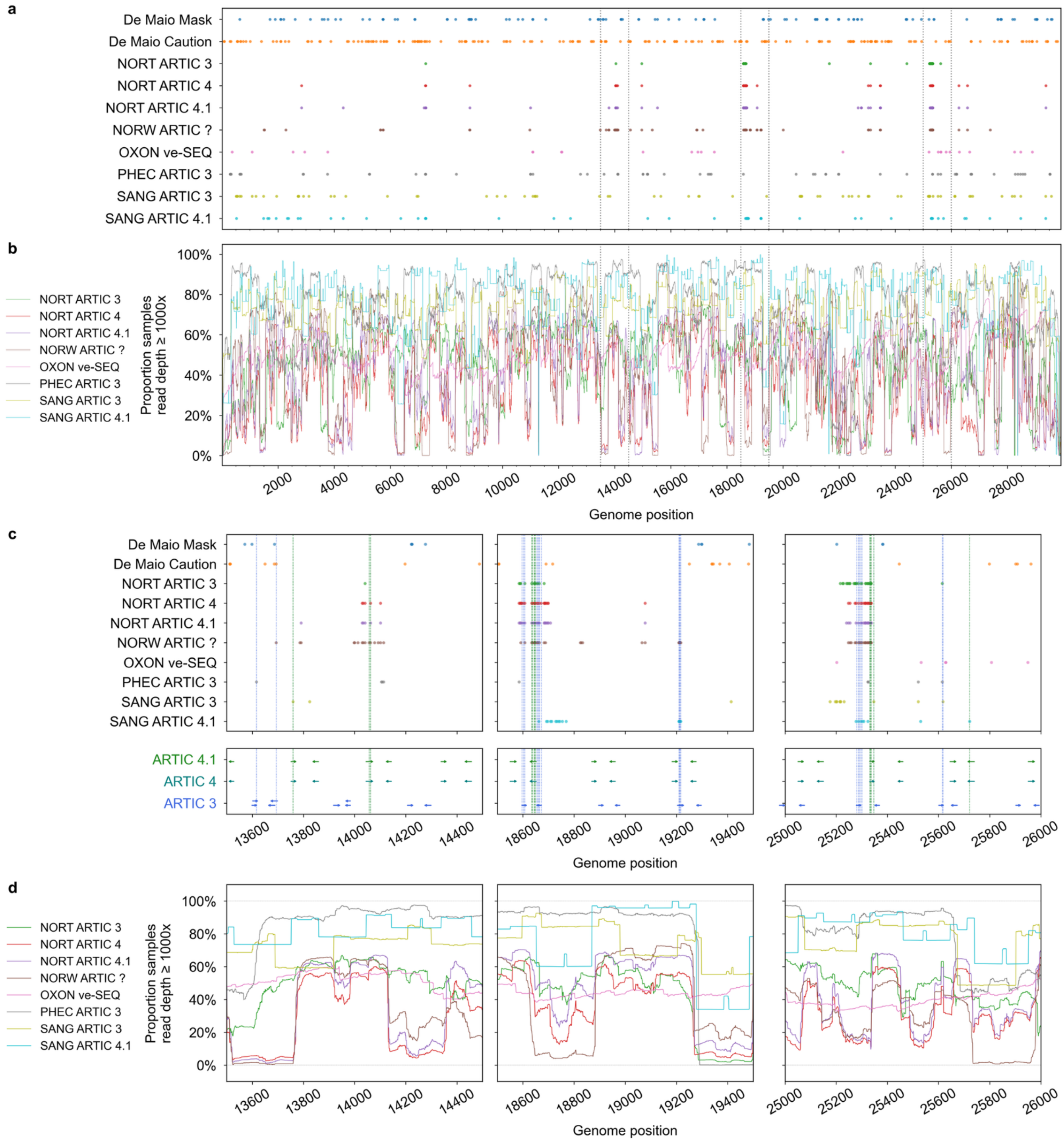


**Supplementary Figure 8:** Distribution of masked positions and read coverage across the SARS-CoV-2 genome for each dataset. **a**. Masked positions shown as coloured points for each sequencing centre/protocol (colours consistent across panels). The De Maio "Mask" and "Caution" (De Maio et al. 2020) positions are also shown. **b**. Proportion of samples reaching total read depth ≥1000× at each genomic position for the same datasets (coloured lines). Vertical dotted lines mark three genomic regions that are displayed at higher resolution in panels (**c–d**). **c**. Expanded views of the masked positions for each dataset for three genomic regions with high clustering of masked positions. The bottom panel show locations of ARTIC primer-binding sites for v3, v4 and v4.1 drawn as arrows (forward = right-pointing; reverse = left-pointing) within these genomic regions. Vertical dotted lines in the corresponding primer colours indicate masked positions that fall within any of these primer regions. **d**. Coverage profiles for the expanded regions.
